# Population genetic analyses of longitudinal vaginal microbiome reveal racioethnic evolutionary dynamics and prevailing positive selection of *Lactobacillus* adhesins

**DOI:** 10.1101/2023.10.11.561685

**Authors:** Xin Wei, Ming-Shian Tsai, Liang Liang, Liuyiqi Jiang, Chia-Jui Hung, Laura Jelliffe-Pawlowski, Larry Rand, Michael Snyder, Chao Jiang

## Abstract

The vaginal microbiome is important for reproductive health and should constantly evolve in response to dynamic host-microbe interactions. The composition of the vaginal microbiome is associated with ethnicity; however, the evolutionary landscape of the vaginal microbiome, especially in the multi-ethnic context, remains under-studied. In this study, we performed a longitudinal evolutionary vaginal microbiome analysis of 351 samples from 35 pregnant women with diverse ethnic backgrounds and validated the main findings in two cohorts totaling 462 samples from 90 multi-ethnic women. Microbiome alpha diversity and community states showed strong ethnic signatures. *Lactobacillaceae* species had a significantly higher nonsynonymous/synonymous mutation ratio (pN/pS) than non-*Lactobacillaceae* species in all ethnicities. In addition, non-*Lactobacillaceae* anaerobic bacteria were enriched in Black and Latino women, with significantly elevated nucleotide diversity and lower pN/pS in Black women. Intriguingly, the *Lactobacillaceae* species had a large repertoire of positively selected genes, including the human mucin-binding and bacterial cell wall anchor genes, which showed independent, recurrent signatures of positive selection across multiple strains, indicating that the host-microbiome interactions directly drive microbial evolution at the molecular interface. Finally, we propose that the evolutionary metrics reflect the environmental niches of adapting microbes. Our study revealed the extensive ethnic signatures in vaginal microbial diversity, composition, community state, and evolutionary dynamics at species and gene levels, highlighting the importance of studying the host-microbiome ecosystem from an evolutionary perspective.

**Highlights:**

- Extensive ethnic signatures in the vaginal microbial diversity, composition, community state, and evolutionary dynamics were demonstrated.
- Healthy *Lactobacillaceae* species showed lower nucleotide diversity but much more relaxed or even positive selection of genes than non-*Lactobacillaceae* species.
- In Black women, non-*Lactobacillaceae* species displayed higher nucleotide diversity and more stringent negative selection.
- *Lactobacillus* mucin-binding and cell wall anchor genes showed convergent signatures of positive selection across vaginal microbiomes.

## Introduction

Human microbiomes are inextricably linked to human health ^1^. The vaginal microbiome is crucial for women’s gynecological and reproductive health ^2–4^. In contrast to the gut microbiota, vaginal microbiota is characterized by lower species diversity and is frequently dominated by *Lactobacillus* species ^5^. The vaginal microbiota is spatiotemporally dynamic, as influenced by several factors, including internal genetic and physiological factors, such as changes in hormone levels and immunity during pubertal development, menstruation, pregnancy, menopause, and disease ^5–9^. External exposures, or the exposome, such as personal hygiene, sexual behavior, and antibiotic treatment, can also disturb the vaginal microbiome ^10–12^. Ethnicity, a complex factor related to both internal and external factors, can also impact the vaginal microbiome ^13^.

Over the past decade, a growing body of multi-ethnic research has suggested an association between ethnicity and vaginal microbiome ^13–16^. The distribution of community state types (CSTs) varies by ethnicity, with Black and Latino populations having a higher proportion of CST IV than White and Asian populations ^14^. In addition, Black women have a vaginal microbiome with higher alpha diversity and a lower abundance of *Lactobacillus* species, a possible risk factor for spontaneous preterm birth ^16,17^. A Human Microbiome Project (HMP) study found that Black women had an increased prevalence of *Lactobacillus* species in their vaginal microbiome during pregnancy compared with other ethnicities ^13^. These findings suggest that ethnicity is a crucial factor in human vaginal microbiome studies.

Recent microbiome studies have increasingly focused on microbiome intraspecific analyses ^18–20^. Several SNP-based tools have been developed and applied to metagenomic data from multiple sources, including the gut, vagina, and natural environment ^21–26^. Recent studies of the vaginal microbiome have investigated strain diversity, persistence, and vertical inheritance ^20,27–29^. However, few studies have explored the evolutionary dynamics of vaginal microbes and their associations with host characteristics throughout human pregnancy.

The host-microbe interactions shape the vaginal environment ^12,30,31^. Host mucus flow and adhesion changes may select for and against specific microbes on the vaginal wall ^32^. In return, bacteria may regulate the expression of host adhesive molecules and affect the immunity of the host ^33,34^. Since genetic drift plays a minor role in microbial evolution, host factors should impact vaginal microbes as selective pressures ^35^. Previous studies on the gut microbiome have shed light on its evolutionary dynamics. One study associated microbial genetic diversity with family, country, geography, and phylogeography ^36^. Another study found that the SNP distribution of *Bacteroides coprocola* was significantly different in type 2 diabetes patients than in the healthy population ^37^. Recently, Liao *et al.* found that preterm birth may be associated with higher nucleotide diversity of the *Gardnerella* species during pregnancy. Understanding how microbes adapt to the host environment and how microbiome adaptations impact the host can provide new mechanistic insights into the vaginal microbiome-host interactions ^38,39^.

Herein, we analyzed 813 longitudinal vaginal microbiome samples from 125 multi-ethnic women in three independent cohorts. The cohorts included newly acquired 351 samples from 35 pregnant women and 462 samples of 90 women from pregnancy and non-pregnancy validation cohorts ^15^. We hypothesized that ethnicity and other host attributes directly interfere with the evolutionary dynamics of the vaginal microbiome. We explored the vaginal microbiome evolutionary dynamics at the community, species, and genetic levels under diverse ethnic backgrounds. We revealed microbiome-wide racioethnic differential evolutionary dynamics, with signatures of positive selection observed in *Lactobacillus* adhesins, which are instrumental in the direct human-microbiome interface, demonstrating the highly selective nature of host-microbe interactions with broad implications in human health.

## Results

### The vaginal microbiome showed ethnic differences in alpha diversity

We developed an effective host-removal pipeline that increased the non-human reads percentage from 4.5% to 28.7% (Figures S1A and B; Methods) ^40^. A total of 485.2 giga base-pairs (Gbp) of metagenomic data after host reads removal were generated from 351 samples longitudinally collected from 35 ethnically diverse women during pregnancy (Methods, some samples with missing metadata; Figure 1A and Table S1). Participants included Black or African American (N = 9; 100 samples), White or Caucasian (N = 9; 80 samples), Latino (N = 8; 73 samples), Asian (N = 6; 43 samples), Pacific Islander (N = 2; 13 samples), and one missing ethnic information (10 samples). Each participant had an average of 9.1 samples collected. Most samples (90.6%) were collected during the second and third trimesters. Besides ethnicity, other characteristics such as gestational weeks, age, BMI, employment, and housing status were collected (Methods).

**Figure 1.**
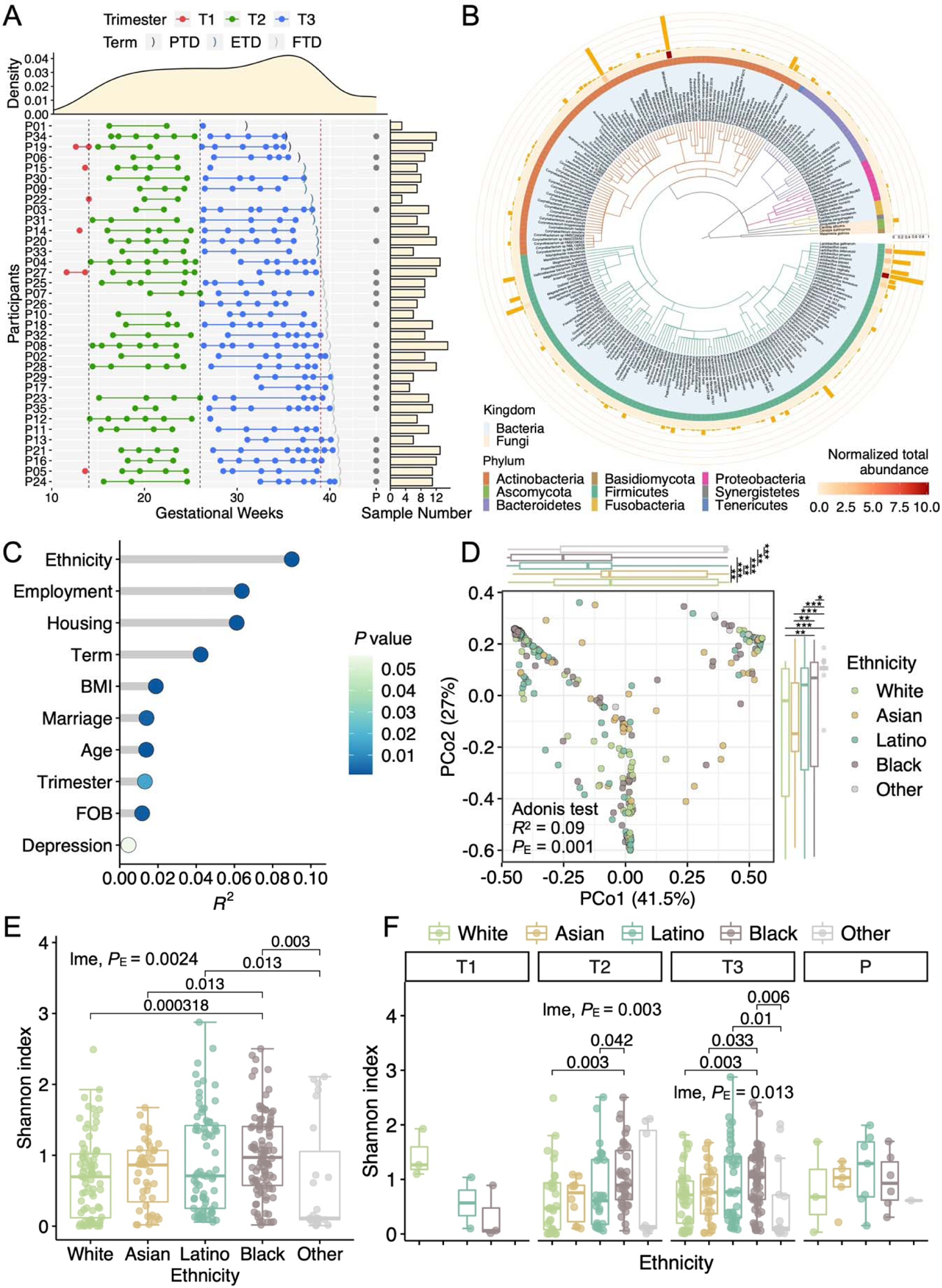
Ethnicity impacts on species diversity of the vaginal microbiome. **(A)** The longitudinal sampling of the vaginal microbiome during and after pregnancy. Each row represents a woman, with bar plots showing the number of samples collected. The points represent the samples collected in the first (T1; red), second (T2; green), third (T3; blue) trimesters, and postpartum (P; grey), respectively. The two black vertical dashed lines divided the three trimesters at weeks 14 and 26 of gestation. The red vertical dashed line marks the full-term delivery (39^th^ week). The brackets represent the delivery time of each woman, with preterm delivery (PTD; <37 weeks), early-term delivery (ETD; 37∼39 weeks), and full-term delivery (FTD; > 39 weeks) colored in light, medium, and dark blue, respectively. The density plot on the top visualizes the distribution of sampling time. **(B)** The phylogenetic tree of the vaginal microbiome in this study. 259 species were identified. The species’ names were colored by the kingdom. The branches and the inner strips were colored by phyla. The outer strip represents the normalized total abundance of each species. The orange bar shows the prevalence of each species. **(C)** The explained variation of several host factors on the beta diversity of the vaginal microbiome. The *P* value and effect size (*R*^2^) were calculated by the multivariate Adonis test. **(D)** The principal coordinates analysis (PCoA) shows the ethnic patterns of the vaginal microbiome. *P*_E_ indicates the *P* value of ethnicity in the multivariate Adonis test. Principal components were compared among ethnicities using the Wilcoxon test. \**P* value < 0.05; *\*\*P* value < 0.01; \*\*\**P* value < 0.001; \*\*\*\**P* value < 0.0001. **(E)** The alpha diversity of different ethnicities. *P* values of significant pair-wise comparisons were shown above the brackets. The ‘lme, *P*_E_’ indicates the *P* value of ethnicity in the multivariate linear mixed-effects (LME) model. **(F)** Alpha diversity was further compared among trimesters in different ethnicities. The LME analysis was performed separately for each trimester.

We parsed the taxonomy using MetaPhlAn4 ^41^. On average, an estimated 83.67% of the metagenomic reads were classified. Two kingdoms, 10 phyla, 119 genera, and 259 species were identified (Figure 1B). The five most prevalent phyla were Firmicutes (100% of 351 samples), Actinobacteria (99.72%), Bacteroidetes (21.65%), the fungal phylum Ascomycota (15.95%), and Proteobacteria (8%). Three fungal species, *Candida albicans*, *Candida dubliniensis*, and *Malassezia globose,* were identified.

To expand the genome database of the vaginal microbiota, we reconstructed 2100 metagenome-assembled genomes (MAGs) of medium quality or better (≥50% completeness and <10% contamination; Methods) ^42^. After clustering the MAGs at 99% average nucleotide identity (ANI), we obtained 424 nonredundant MAGs ^21,43^, of which 240 were high-quality (>90% completeness and <5% contamination, Figure S1C) ^42^. 139 MAGs were further selected as species-level representative genomes (SRGs) at 95% ANI ^44^. All SRGs were identified as bacteria except one archaeon based on the Genome Taxonomy Database (version r207) ^45^. Most bacterial SRGs belong to phylum Firmicutes (including Firmicutes_A/B/C) and Actinobacteria (Figures S1D and E). Notably, a quarter of SRGs (24.46%, 34/139) did not match a reference, defined as unknown SRGs (uSRGs, Figure S1E). To evaluate the relative abundance of the newly recovered genome, we mapped our metagenomic reads to 139 SRGs, and 80.39% of reads were mapped on average (Figure S1F).

We calculated the alpha and beta diversity of the microbiome at the species level to associate with individual characteristics. The Adonis test based on the Bray-Curtis distance showed many factors significantly associated with the vaginal microbiome (*P* < 0.05), including ethnicity, employment, housing, term, BMI, marriage status, age, and whether the father of the baby (FOB) was involved throughout pregnancy (Figure 1C) ^46^. Notably, ethnicity showed the top impact (Figure 1D), followed by employment, housing, and term (Figure S2A-C). Further, we observed significant differences in alpha diversity across ethnic groups (*P*_E_ = 0.00024, *P* value of ‘ethnicity’ in the linear mixed-effects [LME] model), with Black women having the highest Shannon index (Figure 1E), which is also true for the second and third trimesters (*P*_E_ of both LME models < 0.05; Figure 1F). This is consistent with previous findings that the alpha diversity of vaginal microbiome is higher in Black women than in White women ^13,16,17,47^, whereas more ethnic groups were included in our study. Longitudinally, the overall alpha diversity remained stable over gestation time (Figure S2D), consistent with previous findings ^6^, and across ethnic groups (Figure S2E).

### Multi-ethnic pregnant women exhibited divergent vaginal microbiome compositions

We next explored the links between species abundance and ethnicity (Figure S2F and Table S2). Fifteen of the 259 species have been retained after filtering out low-abundance species as described previously (Methods; Figure 2A) ^40^. Seven species were present in more than half of the samples, including *Gardnerella vaginalis* (99.43%), *Lactobacillus crispatus* (96.87%), *Lactobacillus iners* (89.17%), *Fannyhessea vaginae* (88.6%), *Megasphaera* genomosp type 1 (52.42%), *Lactobacillus jensenii* (51.28%), and *Lactobacillus gasseri* (50.43%). Two species, *Actinomycetaceae* SGB989 and *Hungateiclostridiaceae* SGB4003, have never been annotated in the vaginal microbiome. The alpha diversity of samples dominated by *Gardnerella vaginalis* was higher (Figure 2A). *Gardnerella vaginalis* is usually known as the causative agent of bacterial vaginosis (BV) that may increase the risk of preterm birth ^48–50^. Still, its role remains controversial due to its presence in healthy and BV-type vaginal microflora ^51^. *Gardnerella vaginalis* may synergize with other mostly anaerobic bacteria in the vaginal microbiome, such as BV-associated *Fannyhessa vaginae* and *Megasphaera* genomosp type 1 (Figure 2A) ^52–54^. Notably, we observed higher abundance and proportions of *Gardnerella vaginalis* among Black and Latino women in the second and third trimesters (Figure 2B), consistent with previous studies ^13^.

**Figure 2.**
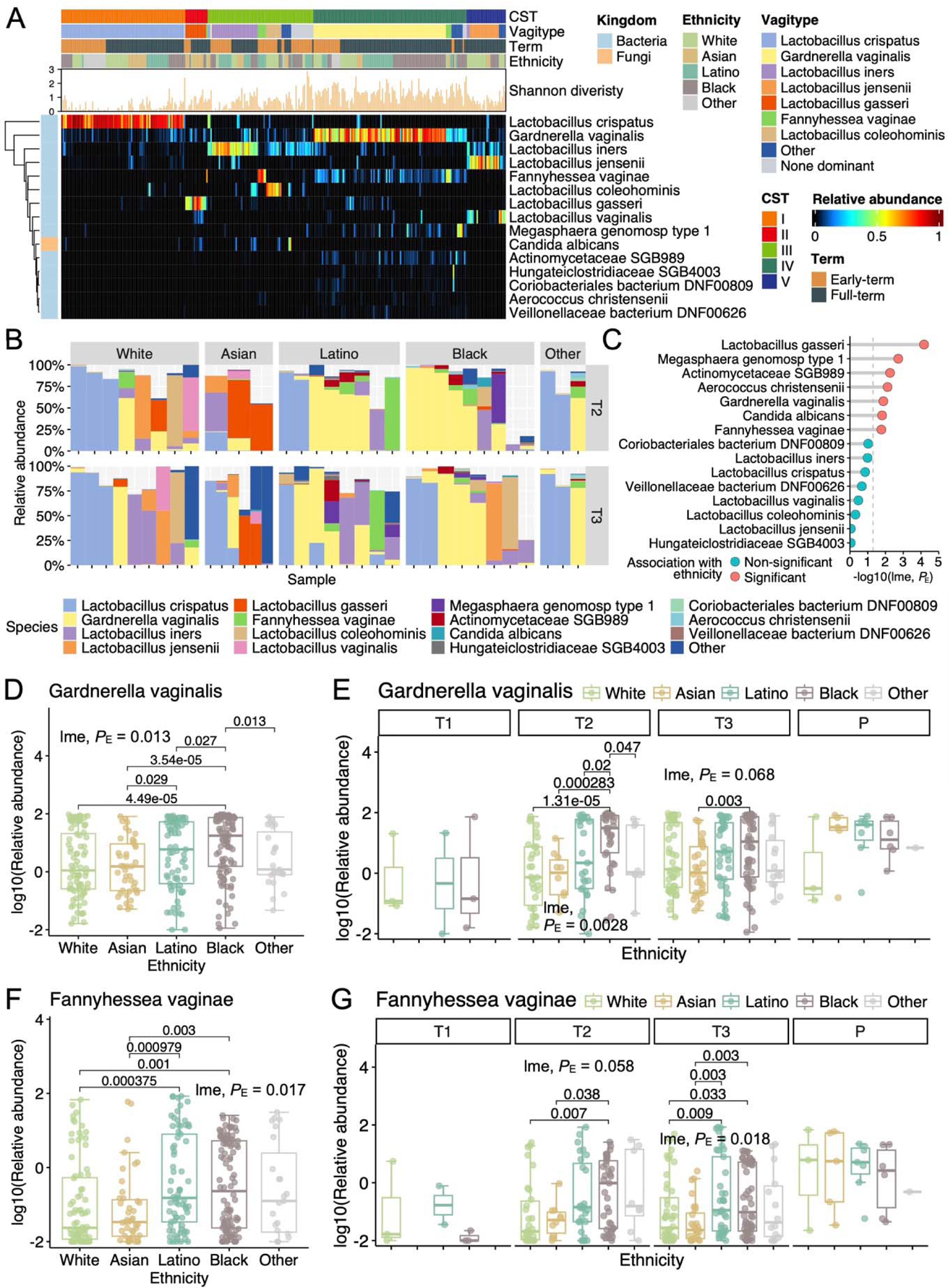
Ethnicity impacts the species composition of the vaginal microbiome. **(A)** The relative abundance of dominant species in the vaginal microbiome. Filter was set as described previously (Methods) ^40^. Common community types (CSTs) or vagitypes were assigned for samples. Early-term group denotes deliveries before the 39^th^ gestational week. **(B)** Vaginal microbiome profiles of 35 pregnant women of White or Caucasian (N = 9), Asian (N = 6), Latino (N = 8), and other ethnicities (N = 3). The samples were rarefied by retaining the earliest sample of the second trimester (T2) and the latest sample of the third trimester (T3) for each woman. **(C)** The associations between ethnicity and the relative abundance of dominant species in the vaginal microbiome. The ‘lme, *P*_E_’ indicates the *P* values of ethnicity in multivariate LME models. A *P* value < 0.05 was considered statistically significant (red). **(D)** The relative abundance of *Gardnerella vaginalis* among ethnicities. **(E)** The relative abundance of *Gardnerella vaginalis* among ethnicities for each trimester. The LME regression was performed separately for each trimester. **(F)** The relative abundance of *Fannyhessea vaginae* among ethnicities. **(G)** The relative abundance of *Fannyhessea vaginae* among ethnicities for each trimester. The LME regression was performed separately for each trimester.

We further analyzed the relationship between microbial abundance and ethnicity using LME models, considering confounding factors. Seven species showed significant associations with ethnicity (Figure 2C), including *Gardnerella vaginalis* and *Fannyhessa vaginae* (*P*_E_ of both LME models < 0.05). Specifically, the prevalence of *Gardnerella vaginalis* was the highest in Black women compared to the other three ethnicities, and it was also higher in Latino than Asian women (Figure 2D). In the second trimester, Black women also had a higher abundance of *Gardnerella vaginalis* than the other three ethnicities (Figure 2E). Compared to White and Asian women, Black and Latino women had a higher abundance of *Fannyhessa vaginae* in all samples (Figure 2F) and the third trimester (Figure 2G). Overall, both *Gardnerella vaginalis* and *Fannyhessa vaginae* had a higher abundance in Black and Latino women than in White and Asian women, at least during the second and third trimesters. Besides, *Lactobacillus gasseri*, *Megasphaera* genomosp type 1, *Actinomycetaceae* SGB989, *Aerococcus christensenii*, and *Candida albicans* also had significant relationships with ethnicity (Figures 2C and S2G-K). Additionally, we observed that Black women seemed to have a lower *Lactobacillus crispatus* than other ethnic groups (Figure S2L). These results suggest that the abundance of most BV-related species and other pathogens is associated with ethnicity ^55^.

### Multi-ethnic vaginal microbiome showed differences between early-term and full-term pregnancies

Preterm birth is an important undesirable outcome, and the correlation between vaginal microbiome and preterm birth has been widely discussed ^4,16,40,56^. To increase statistical power and because early-term delivery (ETD) also had a higher fetal mortality risk ^57,58^, we combined the samples of preterm delivery (PTD) (N = 4) and ETD (N = 9) groups into the ‘early-term’ group (N = 13). The LME regression combining ethnicity and other confounding factors showed no significant difference in the Shannon index between the two groups (*P*_T_ = 0.33, *P* value of ‘term’ in LME model), although a direct Wilcoxon test showed a significantly higher Shannon index in the full-term group (Wilcoxon, *P* = 0.00068; Figure S3A). When modeling separately by trimester, the LME regression of the third trimester showed a significant association between the term and Shannon index with the same trend (LME, *P*_T_ = 0.03; Figure S3B). We also found that the alpha diversity decreased from the second to the third trimester in the ‘early-term’ group (LME, *P*_Trimester_ = 0.053; Figure S3C), contributing to the variation of the alpha diversity over gestation time (Figure S3D), consistent with the previous findings ^56,59^.

We then investigated alpha diversity and early-term birth in conjunction with ethnicity. The LME regressions are insignificant among ethnicities, although the ‘full-term’ Black women had higher alpha diversity than the ‘early-term’ (Wilcoxon, *P* = 0.00057; Figure S3E). Notably, among ‘full-term’ women, Black women had the highest alpha diversity compared to other ethnic groups (LME, *P*_E_ = 0.04; Figure S3F). Thus, although the comparisons related to term have yet to be confirmed by other studies with similar population structures, the differences in alpha diversity among ethnic groups are robust.

We also analyzed the relationship between species abundance and early-term birth, which was strongly influenced by confounding factors (Figure S3G). According to the LME models, we did not observe a significant relationship between early-term birth and the abundance of the three most prevalent species, *Lactobacillus crispatus*, *Gardnerella vaginalis,* and *Lactobacillus iners* (Figures S3G and H-J). However, we found that the three BV-related species or pathogens, *Fannyhessa vaginae*, *Megasphaera* genomosp type 1, and *Aerococcus christensenii*, had a significant relationship with early-term birth (all *P*_T_ of LME < 0.05; Figures S3G and K-M). Specifically, the abundance of *Fannyhessa vaginae* was higher in the ‘early-term’ group than in the ‘full-term’ group in the second (LME, *P*_T_ = 0.0026) and third trimester (LME, *P*_T_ = 0.007, Figure S3K). In addition, *Fannyhessa vaginae* exhibited a decreasing abundance in both ‘early-term’ and ‘full-term’ groups during pregnancy (Figure S3N), similar to the findings of the previous study ^40^.

Besides ethnicity and term, other host factors were linked to the vaginal microbiome (Figures S3O-R). Interestingly, women with part-time employment exhibited higher alpha diversity than employed and unemployed women (Figure S3O). Moreover, unmarried women had a higher relative abundance of *Fannyhessea vaginae* than married women (Figure S3Q). Women with depression had more abundant *Gardnerella vaginalis* (Figure S3R). These results highlight the importance of our diverse population cohort and the inclusion of diverse demographic factors in vaginal microbiome investigation.

### The vaginal community states and transition networks in a multi-ethnic context

We performed CST and vagitype classifications of microbiota based on species abundance profiles (Table S3; Methods) ^6,40^. We observed the consensus clusters between the two classification methods (Chi-square test, *P* < 2.2e-16). Specifically, the dominant species of CST I-III and V correspond to vagitypes *Lactobacillus crispatus*, *Lactobacillus gasseri*, *Lactobacillus iners*, and *Lactobacillus jensenii*, and most communities assigned to vagitype *Gardnerella vaginalis* were also assigned to CST IV (Figure 2A and Table S4).

Principal coordinate analysis (PCoA) showed that the three most distinct groups correspond to CST IV or vagitype *Gardnerella vaginalis*, I or *Lactobacillus crispatus*, and III or *Lactobacillus iners*, respectively (Figures 3A and S4A). The Adonis tests showed CST (*R*^2^ = 0.68) and vagitype (*R*^2^ = 0.78) had much larger effect sizes than other variables in Figure 1C. Meanwhile, we found that alpha diversity was the lowest in CST I and vagitype *Lactobacillus crispatus* but much higher in CST IV and vagitypes *Gardnerella vaginalis,* ‘none dominant’, and the ‘other’ (Figures 3B and S4B).

**Figure 3.**
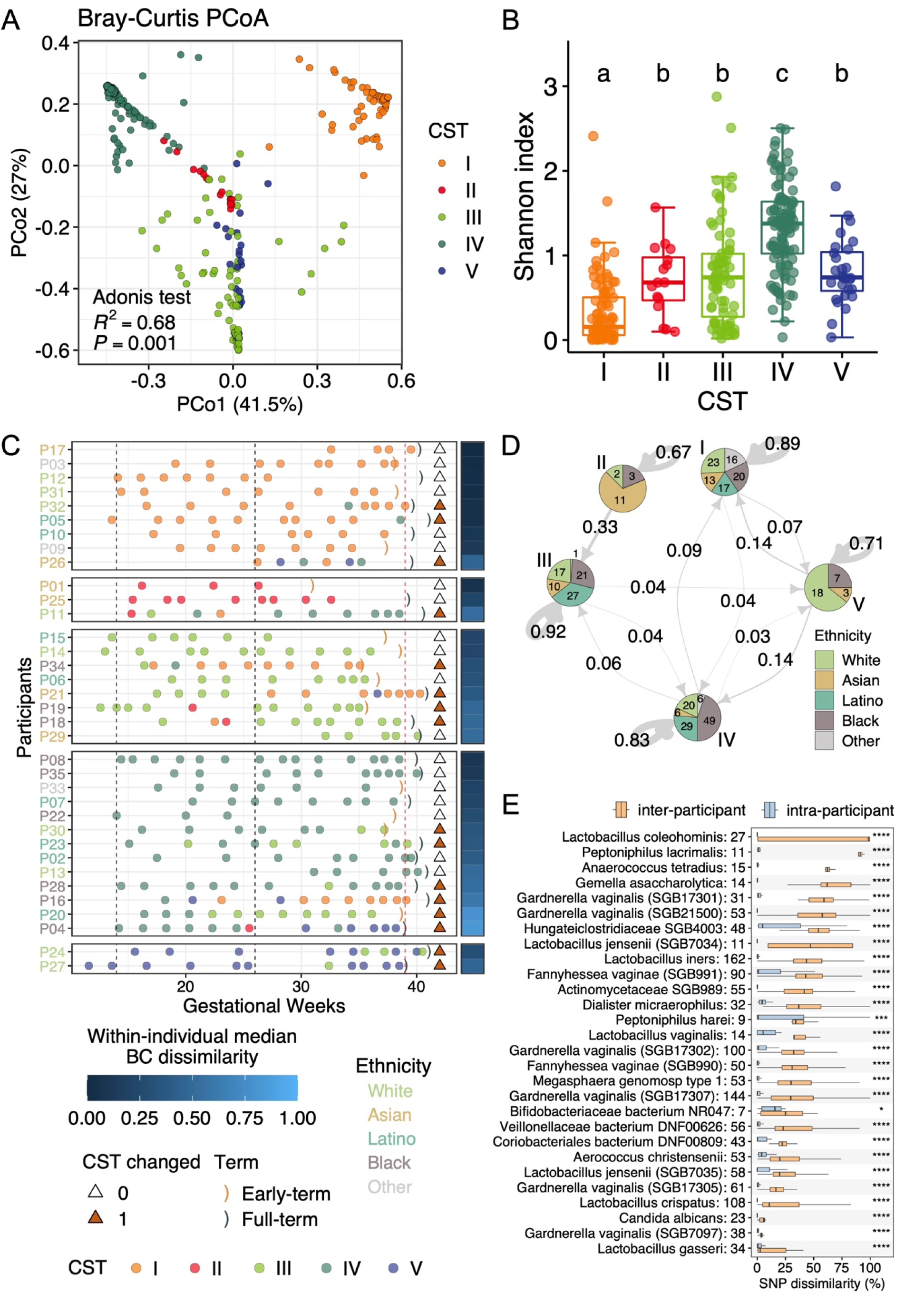
The CSTs showed distinct microbial diversity and ethnic differences and transitioned during pregnancy. **(A)** PCoA analysis shows distinct patterns among vaginal CSTs. **(B)** The alpha diversity of vaginal CSTs. The letters on the top refer to the results of Tukey’s test of all pairwise comparisons. Different letters indicate statistical significance. **(C)** Longitudinal transitions of vaginal CSTs. The triangles indicate whether CSTs changed during pregnancy (filled brown if changed). The vertical dashed lines and brackets are described in Figure 1A. Participants were divided according to the CST of their first samples and were ordered by the median of Bray-Crutis (BC) dissimilarity within individuals. The colors of PIDs indicate ethnicity. **(D)** Transition network of the vaginal CST during pregnancy. The transition probabilities are shown on the edges. Each CST was represented as a pie showing the ethnic composition. The numbers on the pie plots indicate the number of samples. **(E)** Single-nucleotide polymorphism (SNP) haplotype comparisons among different individuals (orange) or the same individuals (blue). Significance indicated by an asterisk (*) was calculated using the Wilcoxon test. References with the same species name are tagged in brackets with the genome ID (SGB+numbers). Numbers following species names indicate the number of applicable samples.

When the CST of the first sample of each participant is assigned as their individual CST, most Black and Latino women (2/3 and 1/2) belonged to CST IV (Table S5), consistent with the previous study ^14^. The vagitype *Gardnerella vaginalis* had the highest proportion in these two ethnic groups (5/9 and 1/2; Table S6), consistent with *Gardnerella vaginalis* being more abundant in Black and Latino women (Figures 2B and D). One-third of Black and Asian women and a quarter of Latino women were assigned CST III (Table S5). The dominant species of CST III is *Lactobacillus iners*, which is associated with BV ^60^. In addition, CST III accounted for 3/4 of PTD women (Table S7) and vagitypes *Lactobacillus iners*, *Fannyhessea vaginae*, and *Megasphaera* genomosp type 1 each accounted for 1/4 of PTD women (Table S8).

We then analyzed the CSTs and vagitypes transition network (Figures 3C and S4C). 45.71% and 65.71% of participants had undergone CST and vagitype transitions during pregnancy (Figure 3C). We depicted vaginal CST/vagitype dynamics as Markov chains (Figure 3D and S4D), generated by inferring the biweekly transition probabilities of CST or vagitype (Tables S9 and 10). According to the self-transition probability, CST II was the most unstable type, with III being the most stable (Figure 3D). The surprisingly high stability of CST IV was probably because up to 86.3% of the CST IV communities belonged to vagitype *Gardnerella vaginalis* (Table S4), which had an 80% self-transition probability (Figure S4D and Table S7). The ‘other’ group and vagitype ‘none dominant’ had the lowest self-transition probability (40% and 50%), suggesting their inherent instability. Interestingly, the self-transition probability of vagitype *Lactobacillus iners* is also low (56.3%), possibly corresponding to the evidence that *Lactobacillus iners* is dominant when the flora is in a transitional stage between abnormal and normal ^61,62^. For inter-CST transition patterns, CST IV had the most bidirectional transitions with other CSTs except for II, similar to DiGiulio *et al*.’s result of the weekly transition ^6^.

Prevalent species not only show differences in relative abundance but also exhibit strain-level variations ^20^. We used StrainPhlAn4 to identify SNP haplotypes in 28 species (see Method). We observed that within-individual genetic differences in SNP haplotypes were significantly lower than the differences between different individuals (Figure 3E). This is consistent with the fact that strains in human microbiome may constitute personal microbial fingerprints ^19,27,63^. Furthermore, we defined distinct haplotype clusters with >70% genetic dissimilarity as ‘strains’ and observed longitudinal strain replacements in seven species/subspecies (Figure S4E).

### Population genetics analyses uncover the distinct ethnic evolutionary dynamics of *Lactobacillaceae* and non-*Lactobacillaceae* species

Next, we conducted an in-depth analysis of the evolutionary dynamics. We profiled the microdiversity for 351 samples with inStrain using 237 NCBI reference genomes or 139 SRGs/MAGs (Methods) ^21^. Evolutionary metrics, including nucleotide diversity (π), recombination rate (1 - linkage disequilibrium [D’]), and the ratio of nonsynonymous to synonymous mutations (pN/pS), were assessed for each genome (Figure 4A; Table S11 and 12). These metrics capture the *in situ* evolutionary dynamics of the vaginal microbiome. The results between corresponding NCBI reference genomes and MAGs are highly consistent (Figure S5A). We henceforth mainly present the results based on NCBI reference genomes for the following reasons: (1) MAGs are rarely complete and may contain chimeric regions ^64,65^; (2) NCBI reference genomes can support multi-cohort analyses. Fifteen genomes were included in the detailed analysis because they had ≥5x coverage across ≥10% of the genome. In total, we detected 29.05 million SNPs in the vaginal microbiome.

**Figure 4.**
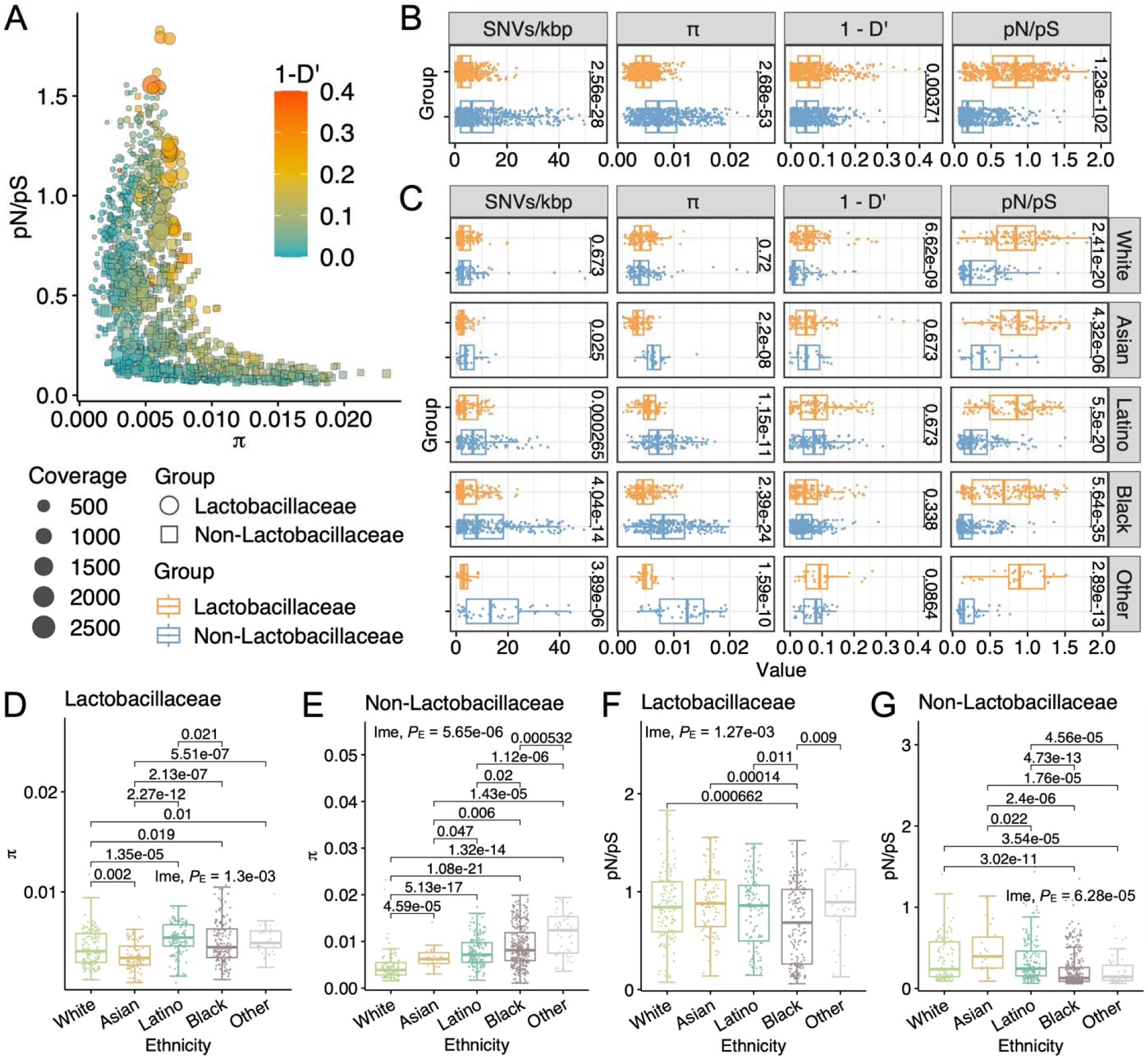
Evolutionary dynamics of *Lactobacillaceae* and non-*Lactobacillaceae* species had distinct ethnic signatures. **(A)** Overview of the evolutionary metrics of 15 species across all samples. The 15 species (displayed in Figure S5B) had sufficient coverage (≥ 5x across ≥ 10% of genome length) in more than 20 samples. Each point or square represents the genome of a certain species. Totaling 1150 genomes in all samples (Table S11). D’ is a measurement of linkage equilibrium ranging from 0 to 1. A higher 1-D’ indicates a higher recombination rate. π, nucleotide diversity; pN/pS, the ratio of non-synonymous rate divided by synonymous rate. **(B)** Comparisons of evolutionary metrics between *Lactobacillaceae* (orange) and non-*Lactobacillaceae* (blue) groups. **(C)** Comparisons of evolutionary metrics between *Lactobacillaceae* (orange) and non-*Lactobacillaceae* (blue) groups for each ethnicity. (**D**-**G**) Comparisons of π (D and E) and pN/pS (F and G) among ethnicities for *Lactobacillaceae* (orange) and non-*Lactobacillaceae* (blue) groups.

Considering the prominent role of the *Lactobacillaceae* family in maintaining low pH in the vaginal micro-environment, we divided the species into *Lactobacillaceae* and non-*Lactobacillaceae* groups. We observed drastically different evolutionary dynamics between these two groups (Figure 4A), reproducible with MAGs-based analyses (Figure S5A). Specifically, the *Lactobacillaceae* group had significantly lower π, higher recombination rate, and higher pN/pS than the non-*Lactobacillaceae* group (Figure 4B). High rates of recombination between strains could cluster lineages together and also lead to gene-specific sweeps and facilitate gene transfer, whereas low rates of recombination could lead to sympatric speciation or more frequent genome-wide sweeps ^66^. We found that the higher recombination rate of the *Lactobacillaceae* group was mainly contributed by *Lactobacillus crispatus* and *Lactobacillus coleohominis*, while some other *Lactobacillaceae* species had even lower recombination rates than non-*Lactobacillaceae* species (Figure S5B). The pN/pS ratio indicates recent selective pressure on a species ^20^. Species undergoing purifying mutations have lower pN/pS ratios, whereas species with low or altered constraints have higher pN/pS ratios ^20^. We observed that *Lactobacillaceae* species experienced neutral or more positive selection than non-*Lactobacillaceae* species. *Lactobacillus iners* exhibited the lowest pN/pS among *Lactobacillaceae* species (Figure S5B).

We next compared the evolutionary dynamics between *Lactobacillaceae* and non-*Lactobacillaceae* groups for each ethnicity (Figure 4C). Strikingly, the pN/pS of the *Lactobacillaceae* group is higher than that of the non-*Lactobacillaceae* group in all ethnicities (Figure 4C), indicating that the more relaxed or even positive selection pressure of *Lactobacillaceae* species is a universal trait. In the *Lactobacillaceae* group, from highest to lowest π were Latino, Black, White, and Asian (Figure 4D). In the non-*Lactobacillaceae* group, from highest to lowest π were Black, Latino, Asian, and White (Figure 4E). To sum up, the vaginal microbiome of Black and Latino women had a higher π than that of White and Asian women. The relationship between π and ethnicity is more significant in the non-*Lactobacillaceae* group (LME, *P*_E_ = 5.65e-06) than in the *Lactobacillaceae* group (LME, *P*_E_ = 1.3e-03). Despite the higher π, Black women had the lowest pN/pS in both the *Lactobacillaceae* and the non-*Lactobacillaceae* groups (Figures 4F and G). The low pN/pS ratios suggest that nonsynonymous mutations in Black women have been purged by purifying selection over time.

To better understand the implications of evolutionary dynamics, we next analyzed the relationships among evolutionary metrics (Figures S5C-I). We observed that coverage did not show consistent correlations with π for the *Lactobacillaceae* and the non-*Lactobacillaceae* groups (Figure S5C). Both groups didn’t show a significant correlation between coverage and pN/pS, indicating the differences in coverage of different bacterial species did not dictate the pN/pS (Figure S5D). In contrast to the overall negative correlation of π and pN/pS in both *Lactobacillaceae* and non-*Lactobacillaceae* groups (Figure S5G), we found that the pN/pS of the *Lactobacillaceae* group was positively correlated with π in White and Asian women (Figure S5H), suggesting that *Lactobacillus* species were prone to accumulate non-synonymous mutations in these two ethnicities. In addition, we noticed that *Lactobacillus iners* had a negative π and pN/pS correlation, unlike other *Lactobacillus* species (Figure S5I), suggesting the particularity of *Lactobacillus iners* from an evolutionary perspective.

### Evolutionary differences in *Lactobacillaceae* and non-*Lactobacillaceae* groups and among ethnicities are validated in independent cohorts

To confirm the divergent vaginal microbiome evolutionary dynamics of two groups of bacteria, we sought to reproduce the evolutionary analyses in public microbiome datasets (Figure S6). Two longitudinal cohorts were suitable for our validation (Table S13), one with 361 samples from 80 non-pregnancy women (validation cohort I) ^27^ and the other with 101 samples from 10 pregnant women (validation cohort II) ^15^. The sample size of validation cohort I is comparable to our study and has diverse ethnicity records. Although the numbers of Latino (N = 4) and Asian (N = 1) women are limited, Black (N = 25) and White women (N = 10) had good representation.

We observed that the *Lactobacillaceae* and the non-*Lactobacillaceae* groups also exhibit distinct evolutionary patterns in the two validation cohorts (Figures S6A and B). The *Lactobacillaceae* group has a lower π, higher recombination rate, and higher pN/pS than the non-*Lactobacillaceae* group in overall samples. Due to the much higher sequencing depth, sample size, and multi-ethnic population, we henceforth focused on validation cohort I (Table S14). Significantly higher pN/pS in the non-*Lactobacillaceae* group can be seen in all ethnicities except for Asian due to the limited sample size (Figure S6C). For the non-*Lactobacillaceae* group, we also found that Black women had higher π and lower pN/pS than other ethnicities (both LME, *P*_E_ < 0.05), but not necessarily for the non-*Lactobacillaceae* group (both LME, *P*_E_ > 0.05; Figures S6D-G). In addition, we similarly found that π and pN/pS are negatively correlated in all samples (Figures S6H and I).

In conclusion, the vaginal *Lactobacillaceae* and *non-Lactobacillaceae* groups showed distinct evolutionary dynamics across ethnicities, regardless of pregnancy. *Lactobacillaceae* species exhibited lower π but tended to experience more relaxed or positive selection pressure than non-*Lactobacillaceae* species. Non-*Lactobacillaceae* species in Black women had higher π and lower pN/pS than other ethnicities, suggesting the genome diversification of non-*Lactobacillaceae* species in Black women by accumulating synonymous mutations. Although the overall coverage of validation cohort I is lower than our study, resulting in slight differences in evolutionary metrics (Figure S6J), the above conclusions are consistent.

### Host and exposome factors were associated with the evolutionary dynamics of the vaginal microbiome

Host and exposome factors may greatly influence the evolutionary dynamics of diverse species ^20^. We systematically evaluated the associations between evolutionary metrics and host and exposome factors (Methods). We found 75 significant associations, of which 51 were significant in the univariate model, 34 were significant in the multivariate model, and 10 were significant in both models (Figure 5A). Notably, the π and pN/pS of *Gardnerella vaginalis* and *Fannyhessea vaginae* had significant associations with ethnicity (*P*_E_ of both univariate and multivariate models < 0.05). These two species had higher π and lower pN/pS among Black women (Figures 5B-E), suggesting that they had higher persistence and experienced purifying selection in the vaginal environment of Black women, resulting accumulation of synonymous mutations over time. Similar to *Gardnerella vaginalis* and *Fannyhessea vaginae*, *Lactobacillus iners* also had higher π and lower pN/pS in Black women (Figures 5F and G). This further illustrates that the evolutionary dynamics of *Lactobacillus iners* were more similar to that of these non-optimal species. Interestingly, we observed that the recombination rate of *Lactobacillus crispatus* was significantly associated with ethnicity, being higher in Latino women (Figure S7A).

**Figure 5.**
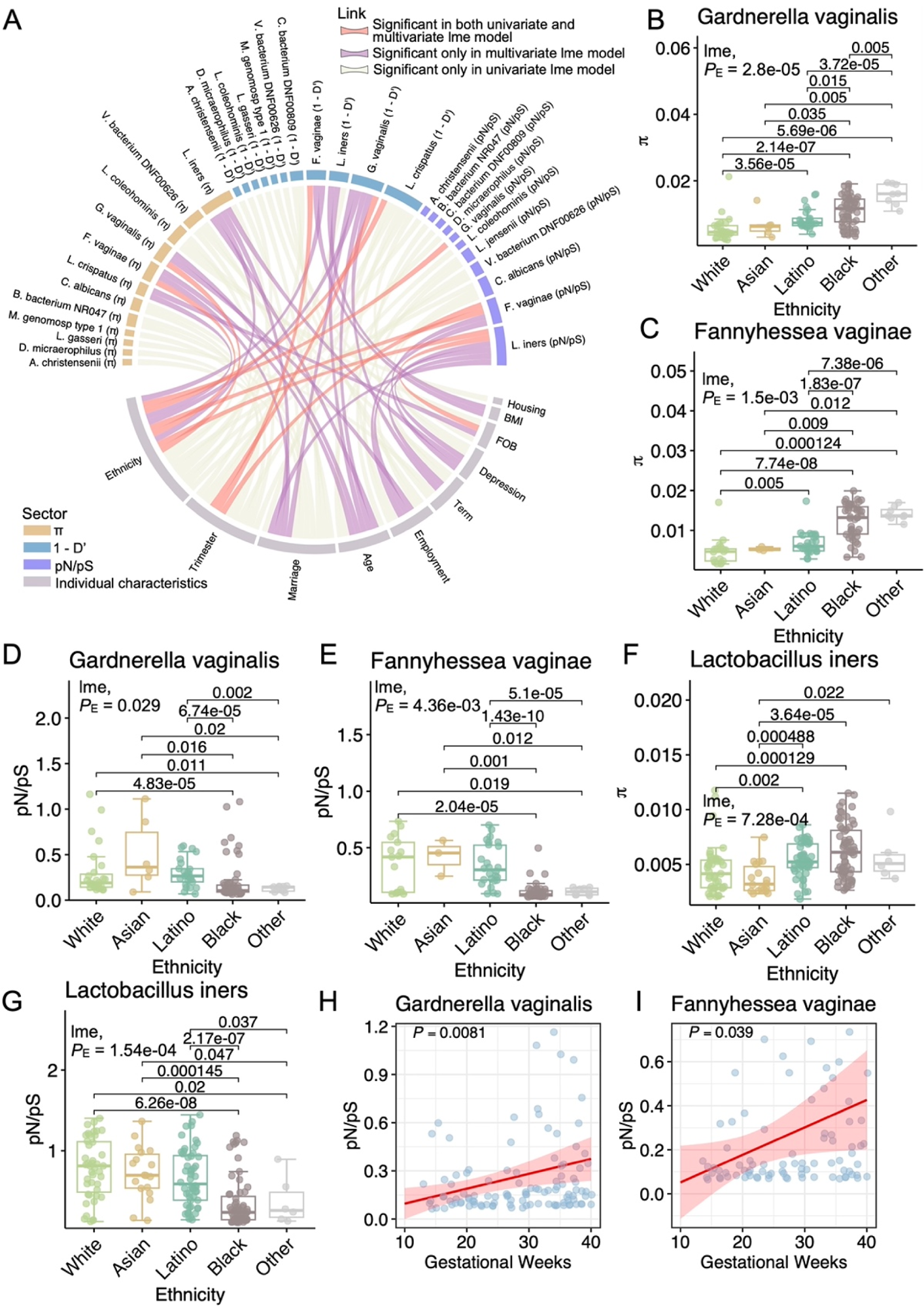
Evolutionary dynamics of the vaginal microbiome were associated with exposome and host factors and changed during pregnancy. **(A)** Overview of 75 significant associations between evolutionary metrics and exposome and host factors. Associations significant in both univariate and multivariate LME models (red lines), the multivariate LME models (purple), and the univariate LME models (light green) were highlighted accordingly. (**B**-**G**) The comparisons of π and pN/pS for several commonly considered non-optimal species among ethnicities. (**H**-**I**) Changes in pN/pS of *Gardnerella vaginalis* and *Fanisella vaginalis* during pregnancy. The fitted line with a 95% confidence interval was shown, calculated based on the regression of the univariate LME model with the participant (subject) as a random effect.

Besides ethnicity, other exposome and host factors were also significantly associated with evolutionary metrics, including trimester, FOB, depression, marriage, and employment (Figures 5A and S7B-L). Trimester had an impact on the evolution of *Gardnerella vaginalis* and *Fannyhessea vaginae*, as their recombination rates and pN/pS increased from the second to the third trimester (Figures S7B-D), corresponding to the increasing trend of pN/pS during pregnancy (Figures 5H and I). Depression was also associated with the evolution of the vaginal microbiome, with higher pN/pS for *Fannyhessea vaginae* and higher π for *Lactobacillus iners* in depressed women (Figures S7G and H). Compared to married women, unmarried women had higher π for *Fannyhessea vaginae* and a higher recombination rate for *Gardneralla vaginalis* but lower pN/pS for *Lactobacillus iners* (Figures S7I-K).

### Deep evolutionary scanning of microbial genes reveals the prevailing positive selection of *Lactobacillus* adhesins

The ability of populations to undergo adaptive evolution depends upon the strength of selection on functional genes to confer competitive advantages. To explore the functional evolutionary propensity of the vaginal microbiome, we performed evolutionary analyses at the gene scale (Methods; Table S15). Genes that encode bacterial adhesion components, integrase, toxin-antitoxin system, and transposase had higher levels of π than the genomic average, whereas ribosomal genes had lower π than the genome average (Figure S8A).

To identify genes and functional modules under positive selection pressure, we focused on the pN/pS and dN/dS of genes. As defined in inStrain, pN/pS is calculated based on positions with at least two alleles present (single nucleotide variant, SNV), unrelated to reference genomes. dN/dS is calculated based on fixed mutations (single nucleotide substitution, SNS) relative to reference genomes ^21^. We visualized the distributions of gene dN/dS and pN/pS across species with sufficient coverage in all samples of both cohorts (Methods). We found that pN/pS was generally higher than dN/dS (Figures 6A and B; S8B and C), suggesting that the vaginal microbiome accumulated more non-fixed non-synonymous mutations, possibly in recent evolutionary time.

**Figure 6.**
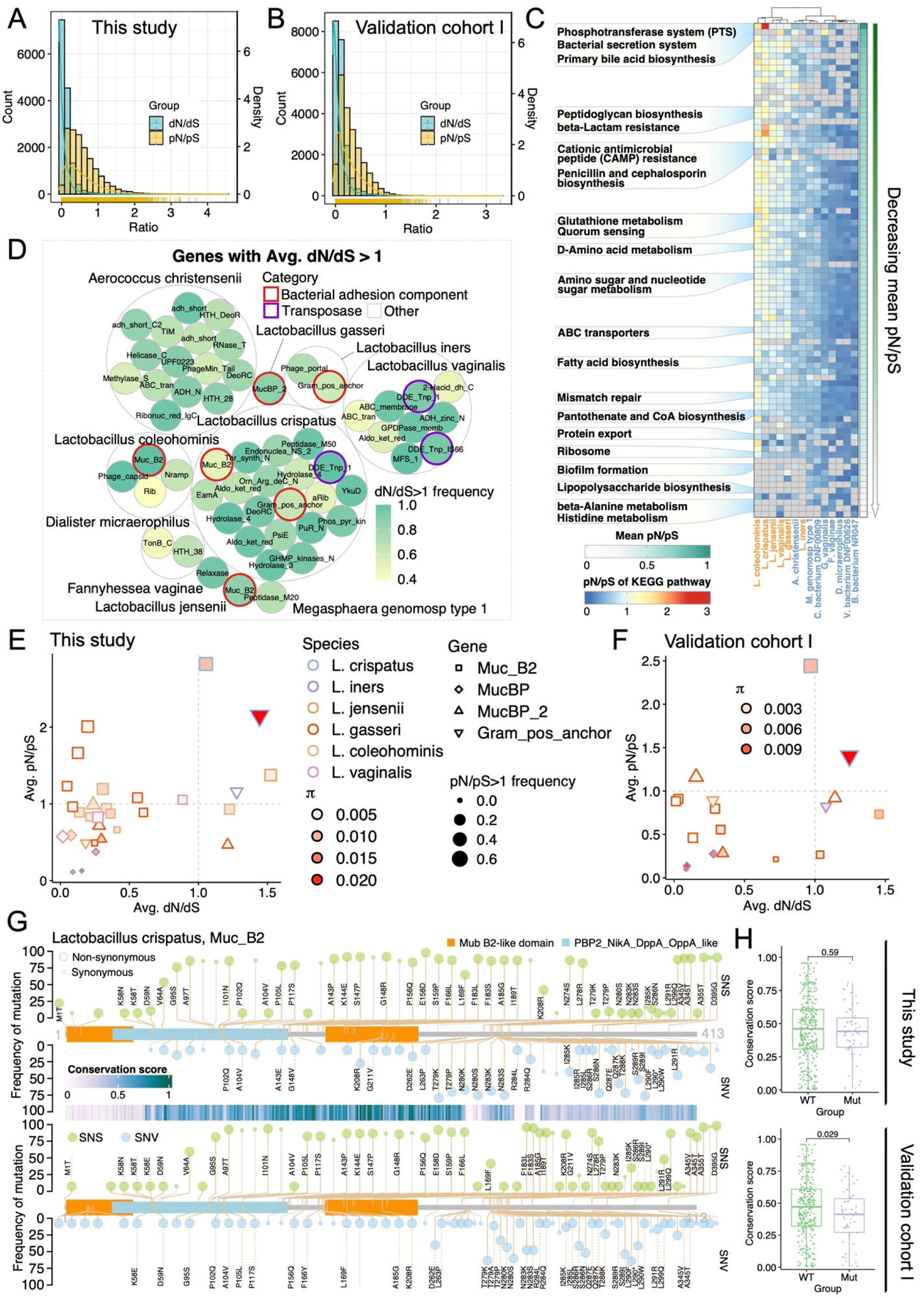
The microbial genes and pathways under positive selections of vaginal microbiome and the mutation hotspots of *Lactobacillus* adhesins. (**A-B**) Distributions of average dN/dS (blue) and pN/pS (yellow) of genes with sufficient coverage (≥ 5x across ≥ 50% of gene length) in more than 10 samples in this study and the validation cohort I. **(C)** The pN/pS of KEGG pathways present in at least three species. The pathways were ordered by the decreasing average pN/pS vertically. *Lactobacillaceae* and non-*Lactobacillaceae* species were colored yellow and blue. Genes that are absent or do not have pN/pS values in some species are filled with gray. **(D)** The genes with average dN/dS greater than 1. Each small circle represents a gene filled based on dN/dS > 1 frequency across samples. 56 genes from 10 species are shown. A larger gray circle packed genes of the same species. Genes of interest were marked by bold circles colored with the assigned categories. (**E**-**F**) The distributions of average dN/dS and pN/pS of *Lactobacillus* adhesins in this study (E) and validation cohort I (F). Genes and species were indicated by the shapes and colors of the points, respectively. The color gradient of points represents average π. The size of the points denotes the frequency of pN/pS > 1 among samples. **(G)** The mutation hotspots on the amino acid sequence of the adhesin gene *muc_B2* of *Lactobacillus crispatus* in this study (top) and validation cohort I (bottom). Two types of mutations, single nucleotide substitution (SNS, green) and single nucleotide variant (SNV, blue), were illustrated on the two sides of the x-axis. Non-synonymous mutations were indicated by bigger dots. The labels of mutations were marked if they were shared in two cohorts. Three known conserved domains on *muc_B2* are represented with color blocks. The sequence and more information about *muc_B2* are recorded in the NCBI Protein database (Accession ID: WP_233263929.1). The conservation score of each amino acid was displayed by the horizontal bar in the middle. **(H)** The comparisons of conservation scores between wild-type loci (WT) and mutated loci (Mut) on adhesin gene *muc_B2* of *Lactobacillus crispatus* in this study (top) and the validation cohort I (bottom).

For an overview of the functional preferences of vaginal microbiome evolution, we examined the pN/pS of KEGG pathways presented by the average pN/pS of involved genes (Figure 6C). Pathways contributing to microbial nutrient utilization, colonization, and persistence in certain environments had higher pN/pS in the *Lactobacillaceae* species, including phosphotransferase system (PTS), primary bile acid biosynthesis, peptidoglycan biosynthesis, and beta-lactam resistance ^67–70^. In contrast, pathways essential for maintaining genome stability, microbial growth, and survival had lower pN/pS, including mismatch repair, pantothenate and CoA biosynthesis, ribosome, and lipopolysaccharide biosynthesis ^71–73^. *Lactobacillaceae* species had higher pN/pS than non-*Lactobacillaceae* species across all present pathways, confirming our previous genome-level results (Figures 4B, S6A, and B). Interestingly, the non-*Lactobacillaceae* species, *Aerococcus christensenii*, which belongs to the same order as *Lactobacillaceae* species, was clustered together with *Lactobacilaceae* species, indicating that the evolutionary dynamics of *Lactobacillaceae* species may be shared beyond the family level (Figure 6C).

We focused on specific genes undergoing positive selection, as indicated by an average dN/dS or pN/pS ratio > 1 across all samples (Figures 6D and S8D). Genes encoding major facilitator superfamily (*MFS_1*), ATP-binding cassette transporter (*ABC_tran*), and transposase (*DDE_Tnp_1*/*DDE_tnp_IS66*) had average dN/dS greater than one (Figure 6D). More importantly, genes that encode bacterial adhesion components: mucin-binding genes (*muc_B2*/*mucBP*/*mucBP_2*) and cell wall anchor genes of gram-positive bacteria (*gram_pos_anchor*) were also under positive selection. These genes are essential for the host-microbe interface at the epithelial surfaces throughout human internal space. Mucins or mucin-like proteins are expressed by epithelial cells in the vagina, digestive and respiratory systems, and are the critical player in the host’s mucosal immunity ^74,75^. The mucosal layer is subjected to bacterial adhesion through binding domains, leading to colonization. Mucin-binding proteins are important components of bacterial adhesion to host mucus and are well-characterized among *Lactobacillus* species ^76^. *Gram_pos_anchor* genes contain LPXTG motif, which attaches to the cell wall peptidoglycan, contributing to the bacterial ability to adhere and colonize surfaces. Our analysis showed that multiple mucin-binding genes and *gram_pos_anchor* genes in several *Lactobacillus* species experienced repetitive positive selection pressure in the vaginal microbial environment. This was observed in both pN/pS and dN/dS metrics (Figures S9A-L). In particular, four genes had average pN/pS and dN/dS > 1, including *muc_B2* and *gram_pos_anchor* in *Lactobacillus crispatus*, *muc_B2* in *Lactobacillus jensenii*, and *gram_pos_anchor* in *Lactobacillus iners* (Figure 6E). We confirmed the findings using uniquely mapped reads (Figure S9M), using SRGs as references (Figure S9N), and in two validation cohorts (Figures 6F and S9O; Table S16).

Since the measurements of selective pressure on genes are the consequence of specific mutations, we visualized the location and frequency of the mutation sites of the four genes with both dN/dS and pN/pS ratios > 1 (Figures S10A-F). Intriguingly, many mutations of these genes were shared with validation cohort I, as shown in Figure 6G for the *muc_B2* in *Lactobacillus crispatus*. In addition, homology alignment analysis showed that mutations are more likely to occur at less conserved positions (Methods; Figures 6H and S10G-L). These analyses showed that shared SNSs or SNVs in adhesins of *Lactobacillus* species were prevalent in two independent cohorts, suggesting either convergent evolution in different populations or the intraspecies genomic diversity was preserved through balancing selection.

## Discussion

Our host-removal experimental pipeline, coupled with deep metagenomic analyses, revealed extensive ethnic signatures present in the vaginal microbial diversity, composition, community state, and, most importantly, evolutionary dynamics. We showed that ethnicity had the greatest impact on microbial diversity compared to other host factors. Notably, Black women exhibited significantly higher alpha diversity than other ethnic groups. We also observed ethnic differences in the relative abundance of presumed non-optimal species, including *Gardnerella vaginalis* and *Fannyhessa vaginae*, which were more prevalent in Black and Latino women. Additionally, we observe varied ethnicity distribution among the CSTs of the vaginal microbiome and showed the CST transition network during pregnancy. These findings not only extend previous knowledge but also provide a comprehensive understanding of vaginal microbiome in a more broad and balanced multi-ethnic context.

In-depth evolutionary analyses of our pregnancy cohort, complemented by validation from two published cohorts, allowed us to examine the intriguing dynamics of *Lactobacillaceae* and non-*Lactobacillaceae* species among ethnicities. Specifically, we found that *Lactobacillaceae* species exhibited lower nucleotide diversity but much more relaxed or even positive selection of genes than non-*Lactobacillaceae* species (Figures 4B and S6B). In Black women, non-*Lactobacillaceae* species displayed greater genome diversification with the accumulation of synonymous mutations (Figures 4E and G; S6E and G). Strain-level diversity is indicative of the demographic history of species, including past changes in population size, population structure, and migration between host microbiomes or other environments ^20^. The lower genetic diversity of the *Lactobacillaceae* species is consistent with the results of Tortelli *et al*., who hypothesized that *Lactobacillus* species had a smaller historical population size or more recent colonization during human history ^20^. In contrast, *Gardnerella vaginalis* and *Fannyhessea vaginae* exhibited greater strain divergence, which is consistent with the hypothesis that the large, long-term populations of these two species may be in the human vagina ancestrally. Besides, we observed that non-*Lactobacillaceae* species have more significant variation among ethnicity than *Lactobacillaceae* species. Unlike the trends of the overall cohort or Black population, the π of non-*Lactobacillaceae* species was as low as *Lactobacillaceae* species in White women (Figures 4C and S6C). This may be because non-*Lactobacillaceae* species such as *Gardnerella vaginalis* and *Fannyhessea vaginae* exhibit less persistence in White women than in Black women, as population persistence is key to genomic diversification ^65^. Taking our multi-ethnic study and Tortelli *et al.*’s together, *Gardnerella vaginalis* and *Fannyhessea vaginae* likely diverged before the migration of modern humans out of Africa and were later colonized by *Lactobacillaceae* species, which may coincide with the elevated use of dairy products ^77,78^.

Recent studies have observed that most microbial genes generally have low pN/pS and inferred a broad purifying selection of microbial genes in the gut and marine microbiome ^18,26,79^. Genes undergoing positive selection are rare in theory, yet they usually play important roles in organismal evolution. The vaginal environment provides a unique interface for host-microbiome interactions due to its low pH and less environmental exposure than the gut. We dug into the evolutionary dynamics at the gene level and observed that many *Lactobacillus* genes were under positive selection. The particularity of the vaginal environment may have driven the positive selection of microbial genes. Especially, we noted the prevailing positive selection of *Lactobacillus* mucin-binding genes (*muc_B2*/*mucBP*/*mucBP_2*) and cell wall anchor genes (*gram_pos_anchor*) in the vaginal microbiome (Figures 6E, F, and S9O). Mucin-binding genes directly interact with mucins which are layered on the host epithelial cells, function as the primary barrier, and coordinate with adaptive and nonadaptive immunity systems ^76^. Surface proteins of gram-positive bacteria contain an LPXTG motif that anchors to the bacterial cell wall^80^. Mucin-binding and cell wall anchor genes are fundamental in bacteria’s ability to colonize and infect host tissues. Their drastically elevated non-synonymous mutation rate indicates a constant demand for evolutionary novelty. The prevailing positive selection of *Lactobacillus* adhesins in both pregnancy and non-pregnancy cohorts indicates the active adaptation of the vaginal microbiota to the host, regardless of pregnancies. Previous studies have provided structural and molecular insights into *Lactobacillus* mucus-binding protein for adapting to gastrointestinal mucus ^81,82^, but the evolutionary dynamics remained unknown. The positively selected sites of mucin-binding proteins may serve as therapeutic targets to deter attachments of unwanted species.

The evolutionary dynamics of a microbe are shaped by its environmental niches. Therefore, microbes that share similar environmental niches may also share similar evolutionary dynamics. We propose a model in which the evolving space of microbes can be mapped by a combination of pN/pS and π. For example, we depicted the microbial evolving space in the vaginal environments of our cohort (Figures S10M and N) and validation cohort I (Figure S10O) in two dimensions. We can observe that *Lactobacillaceae* and non-*Lactobacillaceae* species were distributed diagonally in the second and fourth quadrants, respectively (Figures S10M). Besides *Gardnerella vaginalis* and *Fannyhessa vaginae*, other species in the fourth quadrant were also found to be associated with gynecological disease ^83–87^. Particularly, *Lactobacillus iners* and *Aerococcus christensenii* were near the middle of the diagonal, with the former being an inimical *Lactobacillus* species and the latter an uncommon pathogen ^60,88^. Therefore, we hypothesize that the evolutionary dynamics of a specific microbe could be a powerful tool to characterize its impact on human health in the vaginal environment (Figure S10P). Further analyses of microbiomes in the vaginal or other environments to infer niches of unknown microbes could test this hypothesis.

Despite our compelling findings, several limitations apply. The three cohorts comprise participants from the United States, potentially introducing regional genetic and exposure bias. Moreover, the multi-ethnic demographic characteristics of our cohort may explain the absence of higher microbial alpha diversity in preterm labor women, previously observed in studies of predominantly White women ^4,89^, and has already been disputed by several multi-ethnic studies ^56,59,90,91^. Further studies are necessary to examine whether the positive selection and non-synonymous mutations of *Lactobacillus* adhesins in the vagina lead to functional changes in infectivity, nutrient absorption, and interactions with host epithelium and immunity.

In summary, our study has illuminated the evolutionary landscapes of the vaginal microbiome in a diverse population, with distinct ethnic signatures present at the community, species, strain, and genetic levels. The collection of high-quality metagenome-assembled genomes (MAGs) covers ethnic diversity in the vaginal microbiome and presents a valuable resource for future studies. Notably, bacterial adhesins, among numerous other genes, emerged as the evolutionary hotspots at the human-microbe interface in the vaginal microbiome. Understanding the evolutionary dynamics of these genes offers potential means for modulating adhesin-mucus interactions of undesired species on behalf of the gynecological health of women. Our findings open new avenues to enhance the understanding of the intertwined host-microbiome relationships and human health from a microbial evolutionary perspective.

## Acknowledgments

We gratefully acknowledge the participants who contributed to the study and the teams of research coordinators, the sample processors, and clinicians and nurses who assisted with sample collection. This study was funded by the LSI startup fund, the Fundamental Research Funds for the Central Universities, and the Marc and Lynn Benioff Fund (UCSF7027075) through the California Preterm Birth Initiative (PTBi-CA). We thank the sequencing service in the Genomic Sequencing Service Center at Stanford, and analysis was performed with servers provided by the NECHO computing cluster at Life Sciences Institute at Zhejiang University and the Center for High-Performance Computing at Stanford University.

## Author contributions

L.R. and M.S. conceived the study. M.S., L.R., and C.J. secured the funding and provided overall supervision. C.J. supervised the data analysis. L.L. and L.J.P. coordinated the project to collect samples and metadata. M.T. processed all the samples and generated the data with help from C.H.. X.W. performed all data analyses and visualizations with help from M.T. and L.J.. X.W. and C.J. drafted and revised the manuscript with input from all authors.

## Data availability

The de-identified data for this study have been deposited in the European Nucleotide Archive (ENA) at EMBL-EBI under accession number PRJEB64671. Additional metagenomes were downloaded from the ENA under project IDs PRJNA288562 and PRJNA797778.

## Code availability

Code to replicate all analyses is available from https://github.com/lexinwei/VagMicrobiome.

## Methods

### Study population and sampling procedures

Pregnant women presenting to the University of California, San Francisco (UCSF) Medical Center hospital were invited to participate in this study. All study procedures involving human subjects underwent review and received approval from the institutional review board at UCSF (IRB no. 12-09-0702). The cohort of women in the present analysis was enrolled between November 4, 2014, and September 26, 2018. Inclusion criteria included the age of 18 years or older, the ability to perform the study procedures, and the ability to provide informed consent. All women provided written informed consent before completing an enrollment questionnaire and providing biological samples.

Specimens from the vagina were collected from study enrollment until delivery, and one postpartum sample was also collected in some cases, resulting in 351 samples. 319 samples were collected from 35 women, the rest had missing metadata but were still used for analyses when applicable. Women whose delivery time is before 37 weeks of gestation are considered preterm delivery (PTD), between 37 and 39 weeks are considered early-term delivery (ETD), and over 39 weeks are considered full-term delivery (FTD). All clinical specimens were placed immediately after collection at -20°C until transport to the laboratory for storage at -80°C until processing. Information about age, BMI, ethnicity, employment, marriage, housing status, and disease was captured in a detailed questionnaire completed by each study participant upon enrollment.

### Sample processing and whole shotgun metagenomic sequencing

Literature indicates that the human genome comprises at least 90% of the genomic DNA extracted from vaginal specimens ^40^. To deplete the human genome, a QIAamp DNA microbiome kit was applied to each clinical specimen in accordance with the manufacturer’s protocol. Since the concentration of certain proportions of extracted genomic DNA was too low to directly generate the library, an unbiased amplification method was employed by amplifying the purified DNA at 30°C for 8 hours using a REPLI-g Single Cell kit (Qiagen). The amplified DNA was purified using AMPure XP magnetic beads (Beckman Coulter) according to the supplementary protocol provided by Qiagen. The DNA was eluted in 60 uL 1x TE buffer and quantified using a Qubit dsDNA HS assay kit (Invitrogen). DNA libraries were prepared using a KAPA HyperPlus kit (Roche). Briefly, 500 ng of purified DNA was used for fragmentation. The library was amplified with 3 cycles, followed by size selection for fragments within the 300 bp – 500 bp range using AMPure XP magnetic beads. The library was quantified using a fragment analyzer (Agilent). The libraries were multiplexed, with 91-92 samples per lane, and sequenced on a Novaseq 6000 S4 sequencer with a setting of 2 × 150 bp to generate an average depth of coverage of 26 million reads per sample.

### Whole shotgun metagenomic data pre-processing

Raw sequence data were demultiplexed into sample-specific fastq files using bcl2fastq conversion software from Illumina. Adapter residues were trimmed from both the 5′- and the 3′-end of the reads using the AdapterRemoval v2^92^. The sequences were trimmed for quality using MEEPTOOLS (v0.2) ^93^, retaining reads with a minimum read length of 70 bp and MEEP quality score <1. Human reads were identified and removed from each sample by aligning the reads to the hg19 build of the human genome, using the BWA aligner.

### Taxonomic assignment

Taxonomic classification and relative abundance of species in the metagenomes were obtained using MetaPhlAn4 (v4.0.3) with default parameters ^41^. MetaPhlAn4 integrated information of 1.01_JM prokaryotic reference and metagenome-assembled genomes and defined unique marker genes for 26,970 species-level genome bins for more comprehensive metagenomic taxonomic profiling. The phylogenetic tree was generated using phyloT v2 (https://phylot.biobyte.de/) based on the NCBI taxonomy database and visualized by ggtreeExtra R package (v1.6.1) ^94,95^.

### Filtering out low-abundant taxa

For further analyses related to species abundance, we filtered out low-abundant species that failed to meet both abundance criteria made by Fettweis *et al*. ^40^, that is, (1) 5% of the profiles exhibited an abundance of at least 1%, and (2) at least 15% of profiles exhibited an abundance of at least 0.1%. Fifteen of the 259 species have been retained. Two species, *Actinomycetaceae* SGB989 and *Hungateiclostridiaceae* SGB4003, which have no meaningful names and are denoted by their family plus the MAG ID defined by MetaPhlAn4, have never been annotated in the vaginal microbiome before.

### MAG construction and annotation

#### 1. Metagenomic assembly

Reads from each vaginal sample and the concatenated reads from samples of each participant (referred to as ‘coReads’) were assembled using MetaWRAP (v1.3.2) with the options “--metaspades” and “--megahit”, respectively ^96^. The resulting contigs with lengths less than 1000bp were filtered out. Individual assembly results in 1755 contigs on average. The average N50 is 32402 bp. coReads assembly results in 8663 contigs on average. The average N50 is 29277 bp.

#### 2. Binning

The contigs generated by both individual and coReads assembly were binned using MetaWRAP with the options “--metabat2 --concoct --maxbin2”. The resulting bins were refined with the command “metaWRAP -c 50 -x 10 bin_refinement”, which retained the bins with > 50% completeness and < 10% contamination. A total of 2100 bins were generated.

#### 3. Dereplication and species-level clustering

To remove the redundancy of bins, we integrated the bins derived from individual samples and coReads and clustered them using dRep (v3.4.2) with 99% nucleotide identity (ANI) separately for each participant ^97^. The resulting 683 participant-specific genome sets were further clustered by 99% ANI across participants, and 424 bins were retained. The 424 nonredundant MAGs were further clustered into species-level representative genomes (SRGs) at the threshold of 95% ANI, resulting in 139 SRGs.

#### 4. Taxonomic classification

Phylogenetic inference was conducted by GTDB-Tk (v2.0.0) to classify the 424 nonredundant MAGs based on the Genome Taxonomy Database (GTDB r207; https://gtdb.ecogenomic.org/) ^45,98^. To resolve the taxonomic distribution of the vaginal MAGs, phylogenetic trees were inferred from concatenated sets of single-copy marker genes (120 bacterial or 53 archaeal genes). The recovered 424 nonredundant MAGs spanned ten phyla, including bacteria (423 MAGs) belonging to Actinobacteria (246), Firmicutes (including Firmicutes_A/B/C; 156), Bacteroidota (9), and Patescibacteria (8), etc., and one archaeon from the Thermoproteota. SRGs were considered known SRGs if at least one MAG within the same cluster had species-level annotation. Otherwise, the SRGs were considered unknown SRGs (uSRGs).

#### 5. Phylogenetic analysis

Maximum-likelihood trees were generated de novo using the protein sequence alignments produced by GTDB-Tk: we used IQ-TREE (v2.0.3) to build a phylogenetic tree of the 138 bacterial species ^99^. The best-fit model was automatically selected by ‘ModelFinder’ based on the Bayesian information criterion (BIC) score ^100^. The LG+R7 model was chosen for building the bacterial tree. Trees were visualized and annotated with ggtreeExtra (v1.6.1) ^95^.

#### 6. Relative abundance of SRGs

To calculate the relative abundance of SRGs, reads from each sample were mapped to the set of 139 SRGs using CoverM (https://github.com/wwood/CoverM).

### Calculation of alpha diversity

Based on the MetaPhlAn-4 profiles, we calculated the Shannon index using the utility R script ‘calculate_diveristy.R’ provided by MetaPhlAn4, which embeds the alpha() function of the microbiome R package (v1.18.0) to calculate alpha diversity.

### Community state types (CSTs) and vagitypes assignment

CSTs are clusters of community samples with similar compositions of microbial taxa. We referred to the method of DiGiulio *et al.* to define the vaginal CSTs using Bray-Curtis distance ^6^. Specifically, the Bray-Curtis distance among MetaPhlAn4 profiles was calculated using the vegdist() function in the vegan R package (v2.6-4). The resulting distance matrix was ‘denoised’ by extracting the most significant PCoA eigenvectors. The partitioning around medoids algorithm (pam in R) was applied to these PCoA distances. The number of clusters (k=5) was determined from the gap statistic. This clustering effectively separated vaginal communities into five distinct CSTs: four dominated by different *Lactobacillus* species and one CST with greater diversity and without a dominant *Lactobacillus* strain. These CSTs were analogous to those described previously and were named in accordance ^14^.

According to Fettweis *et al.* ^40^, vagitypes were assigned as the taxon with the largest proportion of reads. Samples in which the largest proportion was less than 30% were not assigned a CST/vagitype. Here, these samples were referred to as vagitype ‘none dominant’, possibly having a high unclassified ratio or diverse species with an abundance lower than 30%. The ‘other’ group includes vagitypes defined in no more than five samples.

Both CSTs and vagitypes classifications were applied to clustering the vaginal microbial community based on MetaPhlAn4 profiles. On the one hand, the concise five CST types have been generalized among many studies, which is convenient for comparing studies or populations. Vagitypes, on the other hand, are used as prototypes for vaginal microbiomes and may vary depending on the predominant species of a particular population ^101^.

### Estimating CST/vagitype transition rates

Vaginal communities exhibited inter-state transitions during pregnancy. We represented vaginal CST and vagitype dynamics as a Markov chain, using DiGiulio *et al*.’s method ^6^. The Markov chain was generated by inferring the biweekly transition probabilities of CST or vagitype, considering the distribution of interval times of sequential samples.

### Strain-level SNP haplotypes and strain replacement

Strain SNP haplotypes were generated using StrainPhlAn4 (v4.0.3) ^22,41^. This method can reconstruct the sample-specific genetic makeup of individual strains using unique clade-specific marker genes exploited by MetaPhlAn4. Reconstructed markers with < 80% breadth of coverage were discarded. Consensus sequences were then trimmed by removing the first and last 50 bases because the terminal positions have lower coverage due to the limitations in mapping reads against truncated sequences ^22^. Next, the markers presented in < 80% of samples were removed. Samples with a percentage of markers < 80% were removed from the alignment. Referring to Chen *et al*.’s method ^63^, we used the multiple sequence alignment to generate a phylogenetic distance matrix by applying the Kimura 2-parameter method from the EMBOSS software (v6.6.0)^102^. Finally, we calculated the SNP haplotype differences of the dominant strain in 28 species/subspecies that were present in both inter- and intra-individual sample pairs. To identify distinct strain clusters within species, the SNP haplotype distance matrix was normalized by dividing maximal distance, and hierarchical clustering was performed with the ‘complete’ method by R basic function hcluster(). Strain clusters were defined at a tree height of 0.7 (70% dissimilarity).

Some species exhibited strain replacement during pregnancy. We represented the strain replacements as Markov chains, similar to the visualization of CST/vagitype transition. For species/subspecies with strain replacement, the Markov chain was generated by inferring the transition probabilities of all sequential samples having the SNP haplotype.

### Microdiversity profiling

#### 1. Reads mapping and inStrain parsing

Intrapopulation genetic diversity was characterized by inStrain (v1.6.0) ^21^. A reference genome set of 237 species (identified by MetaPhlAn4) with available NCBI genomes was prepared for inStrain profiling. Reads were mapped to the reference genome set using bowtie2 (v2.4.2). The mapping results were parsed with the ‘profile’ module of inStrain to identify SNP within the mapped reads assigned to the reference genome set. Read pairs with > 95% ANI were used for the analysis, and all nonpaired reads were discarded. Reads were filtered with a MapQ score > 2 to reduce the number of mis-mapped reads in the analyses. Nucleotide sites also had to have coverage of > 5 for SNP calling, and the minimum allele frequency to confirm an SNP equals 0.05. SNP was identified as nonsynonymous, synonymous, or intergenic given the gene annotations generated by Prodigal (v.2.6.3) ^103^.

#### 2. Genome and gene filtering

For further analysis, we retained the genomes with > 0.1 breadth_minCov (meaning that at least 10% of the bases in the genome were covered by at least five reads, as minimal coverage [minCov] was set as 5). Genes with > 0.5 breadth_minCov were considered present.

#### 3. Calculation of evolutionary metrics

Population statistics and other metrics were calculated at both the genome and gene levels from the mappings with the inStrain program:

##### a.#Nucleotide diversity

InStrain calculates the nucleotide diversity at every position along the genome by summing the squared frequency of each base, based on all reads: 1−[(A_frequency_)^2^ + (C_frequency_)^2^ + (T_frequency_)^2^ + (G_frequency_)^2^], and then, averages values across genes/genomes.

##### b.#Recombination rates

inStrain calculates the linkage equilibrium (D’) between divergent sites on the same read pair, i.e., a measure of how likely two divergent sites are inherited together on the same read pairs. At least 20 paired reads were required for linkage calculations. D’ values range from 0 to 1, with higher values indicating tight linkage of alleles and lower values indicating higher recombination rates. To help easily understand and visualize, we employed 1-D’ as the measurement of recombination rates.

##### c.#pN/pS and dN/dS

inStrain discriminates the divergent sites into SNVs (single nucleotide variants) and SNSs (single nucleotide substitutions). SNV is described as a genomic locus with multiple alleles present, and SNS is a single nucleotide change that has fixed. SNV represents more short-term mutations that may be cleared or eventually fixed, while SNS is more representative of potential long-term mutations. To measure the selective pressure act on genes, pN/pS was calculated using the formula: (nonsynonymous SNVs/nonsynonymous sites)/(synonymous SNVs/synonymous sites), and dN/dS was calculated using the formula (nonsynonymous SNSs/nonsynonymous sites)/(synonymous SNSs/synonymous sites). The pN/pS or dN/dS ratio > 1 means the bias is towards non-synonymous mutations, indicating the gene is under positive selection. The ratio < 1 means the bias is towards synonymous mutations, indicating the gene is under purifying selection. The ratio = 1 means that non-synonymous and synonymous mutations are at the rate expected by mutating positions randomly. The estimation of gene pN/pS and dN/dS was implemented by inStrain. Genome-wide pN/pS were calculated by taking SNVs of all genes (> 0.5 breadth_minCov) across a genome.

The above pipeline was also applied to the published cohorts to validate the discovery of our study. We also rerun the above analyses for our cohort using 139 SRGs as the reference set.

### KEGG pathway pN/pS analysis

The genes were functionally annotated using eggNOG-mapper (v2.1.10) against the eggNOG database (v5.0.2), which assigned the genes to KEGG pathways ^104,105^. We referred to the method of Sjöqvist *et al.* for the pathway pN/pS analyses ^106^. Only KEGG pathways that were judged relevant for microbial genomes were included; the pathways belonging to categories 1.1– 1.12, 2.1–2.4, 3.1, 4.4–4.5, and 6.11 (https://www.genome.jp/kegg/pathway.html#genetic). For genes assigned to multiple pathways, all of the assignments were used. The pN/pS of a KEGG pathway was the average pN/pS of involved genes.

### Gene annotation

The protein sequences of genes were searched by hmmscan (HMMER v3.3.2) against the Pfam database (v34.0) with E-value cutoff 1e-5 ^107,108^. More information about *Gram_pos_anchor* of *Lactobacillus iners*, *Muc_B2* and *Gram_pos_anchor* in *Lactobacillus crispatus*, and *Muc_B2* in *Lactobacillus jensenii* were deposited in the NCBI Protein database with the following accession IDs: QFZ99646.1, WP_233263929.1, WP_158181626, and WP_075362409.1. The annotation of functional units in proteins can be accessed by the related link to the Conserved Domain Database (CDD; https://www.ncbi.nlm.nih.gov/cdd).

### Calculation of conservation score

We searched the homologous sequences of genes using the BLASTp. The first 100 hit sequences with the lowest E-values were aligned using the constraint-based multiple alignment tool (COBALT) in the NCBI C++ toolkit ^109^. The conserve() function of the bio3d R package (v2.4-4) was used to calculate the conservation score of every locus of genes based on the alignments of homologous sequences ^110^.

### Visualization of gene mutation hotspots

We used the trackViewer R package (v1.32.1) to visualize the mutations on genes in the form of a lollipop plot ^111^. The mutation was displayed if it was present in more than 1 sample.

### Statistical analysis

#### Principal coordinates analysis (PCoA)

We applied the vegdist() function from the vegan R package (v2.6-4) to calculate the Bray-Curtis dissimilarity based on the species composition given by MetaPhlAn4. Subsequently, classical multidimensional scaling was carried out based on the Bray-Curtis distance matrix to obtain different principal coordinates.

#### Multivariate Adonis test

We performed the permutational multivariate analysis of variance based on the Bray-Curtis (BC) distance matrix by the vegan::adonis2() function with the formula: BC distance matrix ∼ Ethnicity + Employment + Housing + Term + BMI + Marriage + Age + Trimester + FOB + Depression.

#### Microbial associations to host factors

The species diversity, composition, and evolutionary metrics of the vaginal microbiome can be potentially influenced by a variety of factors. The subject factor is the largest source of variation in sampled communities. Therefore, we employed linear mixed-effects (LME) models throughout our study to investigate the microbial associations to host factors. The LME modeling was performed using the lme() function from the nlme R package (v3.1-162). Multivariate LME models incorporated ethnicity, employment, housing status, term, BMI, marriage, age, FOB, depression, trimester, and a random subject effect to model the microbial diversity, relative abundance, and evolutionary metrics of each species in the vaginal microbiome. The associations with LME *P* value < 0.05 were considered statistically significant.

To identify more possible associations between evolutionary metrics and host factors, besides the multivariate LME model, univariate LME models were also employed for the regression with the formula: dependent variable ∼ independent variable + (1| subject). The significant associations in univariate and/or multivariate LME models were visualized using the circlize R package (v0.4.15)^112^. Besides, two-sided Wilcoxon tests and linear regression were applied for categorical and numerical factors.

#### Evaluating trends with gestational time

Progressive changes in microbial diversity, relative abundance, and pN/pS over the course of pregnancy were fitted by univariate LME models, taking gestational week as the fixed effect and participant as the random effect. Additionally, generalized additive mixed models (GAMMs) incorporating ethnicity, employment, housing status, term, BMI, marriage, age, FOB, depression, a smoother for gestational week, and a random subject effect were used to longitudinally model log-transformed Shannon index of vaginal microbiome. Models were fit using the gamm4 R package (v0.2-6).

All statistical analyses were performed using R language and environment (R 2022, http://www.rproject.org) version 4.2.0 and the add-on packages. The code is publicly available: https://github.com/lexinwei/VagMicrobiome.

## References

1. Proctor, L.M., Creasy, H.H., Fettweis, J.M., Lloyd-Price, J., Mahurkar, A., Zhou, W., Buck, G.A., Snyder, M.P., Strauss, J.F., Weinstock, G.M., et al. (2019). The Integrative Human Microbiome Project. Nature 569, 641–648. 10.1038/s41586-019-1238-8.

2. France, M., Alizadeh, M., Brown, S., Ma, B., and Ravel, J. (2022). Towards a deeper understanding of the vaginal microbiota. Nature Microbiology 7, 367–378. 10.1038/s41564-022-01083-2.

3. Ma, B., Forney, L.J., and Ravel, J. (2012). Vaginal Microbiome: Rethinking Health and Disease. Annual Review of Microbiology 66, 371–389. 10.1146/annurev-micro-092611-150157.

4. Freitas, A.C., Bocking, A., Hill, J.E., Money, D.M., Money, D., Bocking, A., Hemmingsen, S., Hill, J., Reid, G., Dumonceaux, T., et al. (2018). Increased richness and diversity of the vaginal microbiota and spontaneous preterm birth. Microbiome 6, 117. 10.1186/s40168-018-0502-8.

5. Greenbaum, S., Greenbaum, G., Moran-Gilad, J., and Weintraub, A.Y. (2019). Ecological dynamics of the vaginal microbiome in relation to health and disease. American Journal of Obstetrics and Gynecology 220, 324–335. 10.1016/j.ajog.2018.11.1089.

6. DiGiulio, D.B., Callahan, B.J., McMurdie, P.J., Costello, E.K., Lyell, D.J., Robaczewska, A., Sun, C.L., Goltsman, D.S.A., Wong, R.J., Shaw, G., et al. (2015). Temporal and spatial variation of the human microbiota during pregnancy. Proceedings of the National Academy of Sciences 112, 11060–11065. 10.1073/pnas.1502875112.

7. Ravel, J., Brotman, R.M., Gajer, P., Ma, B., Nandy, M., Fadrosh, D.W., Sakamoto, J., Koenig, S.S.K., Fu, L., Zhou, X., et al. (2013). Daily temporal dynamics of vaginal microbiota before, during and after episodes of bacterial vaginosis. Microbiome 1, 29. 10.1186/2049-2618-1-29.

8. Gajer, P., Brotman, R.M., Bai, G., Sakamoto, J., Schütte, U.M.E., Zhong, X., Koenig, S.S.K., Fu, L., Ma, Z., Zhou, X., et al. (2012). Temporal Dynamics of the Human Vaginal Microbiota. Science Translational Medicine 4, 132ra152–132ra152. 10.1126/scitranslmed.3003605.

9. Romero, R., Hassan, S.S., Gajer, P., Tarca, A.L., Fadrosh, D.W., Nikita, L., Galuppi, M., Lamont, R.F., Chaemsaithong, P., Miranda, J., et al. (2014). The composition and stability of the vaginal microbiota of normal pregnant women is different from that of non-pregnant women. Microbiome 2, 4. 10.1186/2049-2618-2-4.

10. Jiang, C., Wang, X., Li, X., Inlora, J., Wang, T., Liu, Q., and Snyder, M. (2018). Dynamic Human Environmental Exposome Revealed by Longitudinal Personal Monitoring. Cell 175, 277–291. 10.1016/j.cell.2018.08.060.

11. Wei, X., Huang, Z., Jiang, L., Li, Y., Zhang, X., Leng, Y., and Jiang, C. Charting the landscape of the environmental exposome. iMeta 1, e50. 10.1002/imt2.50.

12. Kwon, M.S., and Lee, H.K. (2022). Host and Microbiome Interplay Shapes the Vaginal Microenvironment. Frontiers in Immunology 13. 10.3389/fimmu.2022.919728.

13. Serrano, M.G., Parikh, H.I., Brooks, J.P., Edwards, D.J., Arodz, T.J., Edupuganti, L., Huang, B., Girerd, P.H., Bokhari, Y.A., Bradley, S.P., et al. (2019). Racioethnic diversity in the dynamics of the vaginal microbiome during pregnancy. Nature Medicine 25, 1001–1011. 10.1038/s41591-019-0465-8.

14. Ravel, J., Gajer, P., Abdo, Z., Schneider, G.M., Koenig, S.S.K., McCulle, S.L., Karlebach, S., Gorle, R., Russell, J., Tacket, C.O., et al. (2011). Vaginal microbiome of reproductive-age women. Proceedings of the National Academy of Sciences 108, 4680–4687. 10.1073/pnas.1002611107.

15. France, M.T., Fu, L., Rutt, L., Yang, H., Humphrys, M.S., Narina, S., Gajer, P.M., Ma, B., Forney, L.J., and Ravel, J. (2022). Insight into the ecology of vaginal bacteria through integrative analyses of metagenomic and metatranscriptomic data. Genome Biology 23, 66. 10.1186/s13059-022-02635-9.

16. Sun, S., Serrano, M.G., Fettweis, J.M., Basta, P., Rosen, E., Ludwig, K., Sorgen, A.A., Blakley, I.C., Wu, M.C., Dole, N., et al. (2022). Race, the Vaginal Microbiome, and Spontaneous Preterm Birth. mSystems 7, e00017–00022. 10.1128/msystems.00017-22.

17. Fettweis, J.M., Brooks, J.P., Serrano, M.G., Sheth, N.U., Girerd, P.H., Edwards, D.J., Strauss, J.F., Consortium, t.V.M., Jefferson, K.K., and Buck, G.A. (2014). Differences in vaginal microbiome in African American women versus women of European ancestry. Microbiology 160, 2272–2282. 10.1099/mic.0.081034-0.

18. Schloissnig, S., Arumugam, M., Sunagawa, S., Mitreva, M., Tap, J., Zhu, A., Waller, A., Mende, D.R., Kultima, J.R., Martin, J., et al. (2013). Genomic variation landscape of the human gut microbiome. Nature 493, 45–50. 10.1038/nature11711.

19. Lloyd-Price, J., Mahurkar, A., Rahnavard, G., Crabtree, J., Orvis, J., Hall, A.B., Brady, A., Creasy, H.H., McCracken, C., Giglio, M.G., et al. (2017). Strains, functions and dynamics in the expanded Human Microbiome Project. Nature 550, 61–66. 10.1038/nature23889.

20. Tortelli, B.A., Lewis, A.L., and Fay, J.C. (2021). The structure and diversity of strain-level variation in vaginal bacteria. Microbial Genomics 7. 10.1099/mgen.0.000543.

21. Olm, M.R., Crits-Christoph, A., Bouma-Gregson, K., Firek, B.A., Morowitz, M.J., and Banfield, J.F. (2021). inStrain profiles population microdiversity from metagenomic data and sensitively detects shared microbial strains. Nature Biotechnology 39, 727–736. 10.1038/s41587-020-00797-0.

22. Truong, D.T., Tett, A., Pasolli, E., Huttenhower, C., and Segata, N. (2017). Microbial strain-level population structure and genetic diversity from metagenomes. Genome Research 27, 626–638. 10.1101/gr.216242.116.

23. Yilmaz, B., Fuhrer, T., Morgenthaler, D., Krupka, N., Wang, D., Spari, D., Candinas, D., Misselwitz, B., Beldi, G., Sauer, U., and Macpherson, A.J. (2022). Plasticity of the adult human small intestinal stoma microbiota. Cell Host & Microbe 30, 1773–1787.e1776. 10.1016/j.chom.2022.10.002.

24. Lou, Y.C., Olm, M.R., Diamond, S., Crits-Christoph, A., Firek, B.A., Baker, R., Morowitz, M.J., and Banfield, J.F. (2021). Infant gut strain persistence is associated with maternal origin, phylogeny, and traits including surface adhesion and iron acquisition. Cell Reports Medicine 2. 10.1016/j.xcrm.2021.100393.

25. Jingqiu, L., Liat, S., Myrna, S., Bin, Z., Gregory, A.B., and Tal, K. (2023). Microdiversity of the Vaginal Microbiome is Associated with Preterm Birth. bioRxiv, 2023.2001.2013.523991. 10.1101/2023.01.13.523991.

26. Dong, X., Peng, Y., Wang, M., Woods, L., Wu, W., Wang, Y., Xiao, X., Li, J., Jia, K., Greening, C., et al. (2023). Evolutionary ecology of microbial populations inhabiting deep sea sediments associated with cold seeps. Nature Communications 14, 1127. 10.1038/s41467-023-36877-3.

27. Goltsman, D.S.A., Sun, C.L., Proctor, D.M., DiGiulio, D.B., Robaczewska, A., Thomas, B.C., Shaw, G.M., Stevenson, D.K., Holmes, S.P., Banfield, J.F., and Relman, D.A. (2018). Metagenomic analysis with strain-level resolution reveals fine-scale variation in the human pregnancy microbiome. Genome Res 28, 1467–1480. 10.1101/gr.236000.118.

28. France, M., Ma, B., and Ravel, J. (2022). Persistence and In Vivo Evolution of Vaginal Bacterial Strains over a Multiyear Time Period. mSystems 7, e0089322. 10.1128/msystems.00893-22.

29. France, M.T., Brown, S.E., Rompalo, A.M., Brotman, R.M., and Ravel, J. (2022). Identification of shared bacterial strains in the vaginal microbiota of related and unrelated reproductive-age mothers and daughters using genome-resolved metagenomics. PLOS ONE 17, e0275908. 10.1371/journal.pone.0275908.

30. Muzny, C.A., Laniewski, P., Schwebke, J.R., and Herbst-Kralovetz, M.M. (2020). Host– vaginal microbiota interactions in the pathogenesis of bacterial vaginosis. Current Opinion in Infectious Diseases 33, 59–65. 10.1097/qco.0000000000000620.

31. Mahajan, G., Doherty, E., To, T., Sutherland, A., Grant, J., Junaid, A., Gulati, A., LoGrande, N., Izadifar, Z., Timilsina, S.S., et al. (2022). Vaginal microbiome-host interactions modeled in a human vagina-on-a-chip. Microbiome 10, 201. 10.1186/s40168-022-01400-1.

32. McLoughlin, K., Schluter, J., Rakoff-Nahoum, S., Smith, Adrian L., and Foster, Kevin R. (2016). Host Selection of Microbiota via Differential Adhesion. Cell Host & Microbe 19, 550–559. 10.1016/j.chom.2016.02.021.

33. Dohrman, A., Miyata, S., Gallup, M., Li, J.-D., Chapelin, C., Coste, A., Escudier, E., Nadel, J., and Basbaum, C. (1998). Mucin gene (MUC 2 and MUC 5AC) upregulation by Gram-positive and Gram-negative bacteria. Biochimica et Biophysica Acta (BBA) - Molecular Basis of Disease 1406, 251–259. 10.1016/S0925-4439(98)00010-6.

34. Anton, L., Ferguson, B., Friedman, E.S., Gerson, K.D., Brown, A.G., and Elovitz, M.A. (2022). Gardnerella vaginalis alters cervicovaginal epithelial cell function through microbe-specific immune responses. Microbiome 10, 119. 10.1186/s40168-022-01317-9.

35. Li, L., and Ma, Z. (2016). Testing the Neutral Theory of Biodiversity with Human Microbiome Datasets. Scientific Reports 6, 31448. 10.1038/srep31448.

36. Hildebrand, F., Gossmann, T.I., Frioux, C., Özkurt, E., Myers, P.N., Ferretti, P., Kuhn, M., Bahram, M., Nielsen, H.B., and Bork, P. (2021). Dispersal strategies shape persistence and evolution of human gut bacteria. Cell Host & Microbe 29, 1167–1176.e1169. 10.1016/j.chom.2021.05.008.

37. Chen, Y., Pouillot, R., S. Burall, L., Strain, E.A., Van Doren, J.M., De Jesus, A.J., Laasri, A., Wang, H., Ali, L., Tatavarthy, A., et al. (2017). Comparative evaluation of direct plating and most probable number for enumeration of low levels of Listeria monocytogenes in naturally contaminated ice cream products. International Journal of Food Microbiology 241, 15–22. 10.1016/j.ijfoodmicro.2016.09.021.

38. Garud, N.R., and Pollard, K.S. (2020). Population Genetics in the Human Microbiome. Trends in Genetics 36, 53–67. 10.1016/j.tig.2019.10.010.

39. Zhang, J., and Knight, R. (2023). Genomic Mutations Within the Host Microbiome: Adaptive Evolution or Purifying Selection. Engineering 20, 96–102. 10.1016/j.eng.2021.11.018.

40. Fettweis, J.M., Serrano, M.G., Brooks, J.P., Edwards, D.J., Girerd, P.H., Parikh, H.I., Huang, B., Arodz, T.J., Edupuganti, L., Glascock, A.L., et al. (2019). The vaginal microbiome and preterm birth. Nature Medicine 25, 1012–1021. 10.1038/s41591-019-0450-2.

41. Blanco-Míguez, A., Beghini, F., Cumbo, F., McIver, L.J., Thompson, K.N., Zolfo, M., Manghi, P., Dubois, L., Huang, K.D., Thomas, A.M., et al. (2023). Extending and improving metagenomic taxonomic profiling with uncharacterized species using MetaPhlAn 4. Nature Biotechnology. 10.1038/s41587-023-01688-w.

42. Bowers, R.M., Kyrpides, N.C., Stepanauskas, R., Harmon-Smith, M., Doud, D., Reddy, T.B.K., Schulz, F., Jarett, J., Rivers, A.R., Eloe-Fadrosh, E.A., et al. (2017). Minimum information about a single amplified genome (MISAG) and a metagenome-assembled genome (MIMAG) of bacteria and archaea. Nature Biotechnology 35, 725–731. 10.1038/nbt.3893.

43. Brooks, B., Olm, M.R., Firek, B.A., Baker, R., Thomas, B.C., Morowitz, M.J., and Banfield, J.F. (2017). Strain-resolved analysis of hospital rooms and infants reveals overlap between the human and room microbiome. Nature Communications 8, 1814. 10.1038/s41467-017-02018-w.

44. Olm, M.R., Crits-Christoph, A., Diamond, S., Lavy, A., Carnevali, P.B.M., and Banfield, J.F. (2020). Consistent Metagenome-Derived Metrics Verify and Delineate Bacterial Species Boundaries. mSystems 5, e00731–00719. 10.1128/mSystems.00731-19.

45. Parks, D.H., Chuvochina, M., Rinke, C., Mussig, A.J., Chaumeil, P.-A., and Hugenholtz, P. (2021). GTDB: an ongoing census of bacterial and archaeal diversity through a phylogenetically consistent, rank normalized and complete genome-based taxonomy. Nucleic Acids Research 50, D785–D794. 10.1093/nar/gkab776.

46. Lee, S.H., Lazebnik, R., Kuper-Sassé, M., and Lazebnik, N. (2016). Is the Presence of the Father of the Baby during First Prenatal Ultrasound Study Visit Associated with Improved Pregnancy Outcomes in Adolescents and Young Adults? International Journal of Pediatrics 2016, 4632628. 10.1155/2016/4632628.

47. Dominguez-Bello, M.G. (2019). Gestational shaping of the maternal vaginal microbiome. Nature Medicine 25, 882–883. 10.1038/s41591-019-0483-6.

48. Chen, X., Lu, Y., Chen, T., and Li, R. (2021). The Female Vaginal Microbiome in Health and Bacterial Vaginosis. Frontiers in Cellular and Infection Microbiology 11. 10.3389/fcimb.2021.631972.

49. Bretelle, F., Rozenberg, P., Pascal, A., Favre, R., Bohec, C., Loundou, A., Senat, M.-V., Aissi, G., Lesavre, N., Brunet, J., et al. (2014). High Atopobium vaginae and Gardnerella vaginalis Vaginal Loads Are Associated With Preterm Birth. Clinical Infectious Diseases 60, 860–867. 10.1093/cid/ciu966.

50. Menard, J.P., Mazouni, C., Salem-Cherif, I., Fenollar, F., Raoult, D., Boubli, L., Gamerre, M., and Bretelle, F. (2010). High Vaginal Concentrations of Atopobium vaginae and Gardnerella vaginalis in Women Undergoing Preterm Labor. Obstetrics & Gynecology 115, 134–140. 10.1097/AOG.0b013e3181c391d7.

51. Janulaitiene, M., Paliulyte, V., Grinceviciene, S., Zakareviciene, J., Vladisauskiene, A., Marcinkute, A., and Pleckaityte, M. (2017). Prevalence and distribution of Gardnerella vaginalis subgroups in women with and without bacterial vaginosis. BMC Infectious Diseases 17, 394. 10.1186/s12879-017-2501-y.

52. Kusters, J.G., Reuland, E.A., Bouter, S., Koenig, P., and Dorigo-Zetsma, J.W. (2015). A multiplex real-time PCR assay for routine diagnosis of bacterial vaginosis. European Journal of Clinical Microbiology & Infectious Diseases 34, 1779–1785. 10.1007/s10096-015-2412-z.

53. Mendling, W., Palmeira-de-Oliveira, A., Biber, S., and Prasauskas, V. (2019). An update on the role of Atopobium vaginae in bacterial vaginosis: what to consider when choosing a treatment? A mini review. Archives of Gynecology and Obstetrics 300, 1–6. 10.1007/s00404-019-05142-8.

54. Roohbakhsh, E., Mojtahedi, A., Roohbakhsh, Z., Khavari-nejad, R.A., and Amirmozafari, N. (2019). Identification of Gardnerella vaginalis and Atopobium vaginae in Women With Bacterial Vaginosis in Northern Iran. Infectious Diseases in Clinical Practice 27, 81–84. 10.1097/IPC.0000000000000691.

55. Alcendor, D.J. (2016). Evaluation of Health Disparity in Bacterial Vaginosis and the Implications for HIV-1 Acquisition in African American Women. American Journal of Reproductive Immunology 76, 99–107. 10.1111/aji.12497.

56. Stout, M.J., Zhou, Y., Wylie, K.M., Tarr, P.I., Macones, G.A., and Tuuli, M.G. (2017). Early pregnancy vaginal microbiome trends and preterm birth. American Journal of Obstetrics and Gynecology 217, 356.e351–356.e318. 10.1016/j.ajog.2017.05.030.

57. Medicine, T.A.C.o.O.a.G.C.o.O.P.S.f.M.-F. (2013). Committee Opinion No 579: Definition of Term Pregnancy. Obstetrics & Gynecology 122.

58. Spong, C.Y. (2013). Defining “Term” Pregnancy: Recommendations From the Defining “Term” Pregnancy Workgroup. JAMA 309, 2445–2446. 10.1001/jama.2013.6235.

59. Haque, M.M., Merchant, M., Kumar, P.N., Dutta, A., and Mande, S.S. (2017). First-trimester vaginal microbiome diversity: A potential indicator of preterm delivery risk. Scientific Reports 7, 16145. 10.1038/s41598-017-16352-y.

60. Zheng, N., Guo, R., Wang, J., Zhou, W., and Ling, Z. (2021). Contribution of Lactobacillus iners to Vaginal Health and Diseases: A Systematic Review. Frontiers in Cellular and Infection Microbiology 11. 10.3389/fcimb.2021.792787.

61. Jakobsson, T., and Forsum, U. (2007). Lactobacillus iners: a Marker of Changes in the Vaginal Flora? Journal of Clinical Microbiology 45, 3145–3145. 10.1128/JCM.00558-07.

62. Nilsen, T., Swedek, I., Lagenaur, L.A., and Parks, T.P. (2020). Novel Selective Inhibition of Lactobacillus iners by Lactobacillus-Derived Bacteriocins. Applied and Environmental Microbiology 86, e01594–01520. 10.1128/AEM.01594-20.

63. Chen, L., Wang, D., Garmaeva, S., Kurilshikov, A., Vich Vila, A., Gacesa, R., Sinha, T., Segal, E., Weersma, R.K., Wijmenga, C., et al. (2021). The long-term genetic stability and individual specificity of the human gut microbiome. Cell 184, 2302–2315.e2312. 10.1016/j.cell.2021.03.024.

64. Eisenhofer, R., Odriozola, I., and Alberdi, A. (2023). Impact of microbial genome completeness on metagenomic functional inference. ISME Communications 3, 12. 10.1038/s43705-023-00221-z.

65. Okazaki, Y., Nakano, S.-i., Toyoda, A., and Tamaki, H. (2022). Long-Read-Resolved, Ecosystem-Wide Exploration of Nucleotide and Structural Microdiversity of Lake Bacterioplankton Genomes. mSystems 7, e00433–00422. 10.1128/msystems.00433-22.

66. Anderson, R.E., Graham, E.D., Huber, J.A., and Tully, B.J. (2022). Microbial Populations Are Shaped by Dispersal and Recombination in a Low Biomass Subseafloor Habitat. mBio 13, e00354–00322. 10.1128/mbio.00354-22.

67. Francl, A.L., Thongaram, T., and Miller, M.J. (2010). The PTS transporters of Lactobacillus gasseri ATCC 33323. BMC Microbiology 10, 77. 10.1186/1471-2180-10-77.

68. Prete, R., Long, S.L., Gallardo, A.L., Gahan, C.G., Corsetti, A., and Joyce, S.A. (2020). Beneficial bile acid metabolism from Lactobacillus plantarum of food origin. Scientific Reports 10, 1165. 10.1038/s41598-020-58069-5.

69. Schneewind, O., and Missiakas, D.M. (2012). Protein secretion and surface display in Gram-positive bacteria. Philosophical Transactions of the Royal Society B: Biological Sciences 367, 1123–1139. 10.1098/rstb.2011.0210.

70. Sirichoat, A., Flórez, A.B., Vázquez, L., Buppasiri, P., Panya, M., Lulitanond, V., and Mayo, B. (2020). Antibiotic Susceptibility Profiles of Lactic Acid Bacteria from the Human Vagina and Genetic Basis of Acquired Resistances. International Journal of Molecular Sciences 21, 2594.

71. Castañeda-García, A., Prieto, A.I., Rodríguez-Beltrán, J., Alonso, N., Cantillon, D., Costas, C., Pérez-Lago, L., Zegeye, E.D., Herranz, M., Plociński, P., et al. (2017). A non-canonical mismatch repair pathway in prokaryotes. Nature Communications 8, 14246. 10.1038/ncomms14246.

72. Spry, C., Kirk, K., and Saliba, K.J. (2008). Coenzyme A biosynthesis: an antimicrobial drug target. FEMS Microbiology Reviews 32, 56–106. 10.1111/j.1574-6976.2007.00093.x.

73. Servais, C., Vassen, V., Verhaeghe, A., Küster, N., Carlier, E., Phégnon, L., Mayard, A., Auberger, N., Vincent, S., and De Bolle, X. (2023). Lipopolysaccharide biosynthesis and traffic in the envelope of the pathogen Brucella abortus. Nature Communications 14, 911. 10.1038/s41467-023-36442-y.

74. Van Tassell, M.L., and Miller, M.J. (2011). Lactobacillus Adhesion to Mucus. Nutrients 3, 613–636. 10.3390/nu3050613.

75. Dharmani, P., Srivastava, V., Kissoon-Singh, V., and Chadee, K. (2008). Role of Intestinal Mucins in Innate Host Defense Mechanisms against Pathogens. Journal of Innate Immunity 1, 123–135. 10.1159/000163037.

76. Devi, S.M., and Halami, P.M. (2017). Diversity and evolutionary aspects of mucin binding (MucBP) domain repeats among Lactobacillus plantarum group strains through comparative genetic analysis. Systematic and Applied Microbiology 40, 237–244. 10.1016/j.syapm.2017.03.005.

77. Rosen, E.M., Martin, C.L., Siega-Riz, A.M., Dole, N., Basta, P.V., Serrano, M., Fettweis, J., Wu, M., Sun, S., Thorp Jr, J.M., et al. (2022). Is prenatal diet associated with the composition of the vaginal microbiome? Paediatric and Perinatal Epidemiology 36, 243–253. 10.1111/ppe.12830.

78. B. Moura, G., G. Silva, M., and Marconi, C. (2023). Milk and Dairy Consumption and Its Relationship With Abundance of Lactobacillus crispatus in the Vaginal Microbiota: Milk Intake and Vaginal Lactobacillus. Journal of Lower Genital Tract Disease 27, 280–285. 10.1097/lgt.0000000000000736.

79. Shenhav, L., and Zeevi, D. (2020). Resource conservation manifests in the genetic code. Science 370, 683–687. 10.1126/science.aaz9642.

80. Perry, A.M., Ton-That, H., Mazmanian, S.K., and Schneewind, O. (2002). Anchoring of Surface Proteins to the Cell Wall of Staphylococcus aureus: III. LIPID II IS AN IN VIVO PEPTIDOGLYCAN SUBSTRATE FOR SORTASE-CATALYZED SURFACE PROTEIN ANCHORING *. Journal of Biological Chemistry 277, 16241–16248. 10.1074/jbc.M109194200.

81. Etzold, S., Kober, O.I., MacKenzie, D.A., Tailford, L.E., Gunning, A.P., Walshaw, J., Hemmings, A.M., and Juge, N. (2014). Structural basis for adaptation of lactobacilli to gastrointestinal mucus. Environmental Microbiology 16, 888–903. 10.1111/1462-2920.12377.

82. Etzold, S., MacKenzie, D.A., Jeffers, F., Walshaw, J., Roos, S., Hemmings, A.M., and Juge, N. (2014). Structural and molecular insights into novel surface-exposed mucus adhesins from Lactobacillus reuteri human strains. Molecular Microbiology 92, 543–556. 10.1111/mmi.12574.

83. Khan, S., Vancuren, S.J., and Hill, J.E. (2021). A Generalist Lifestyle Allows Rare Gardnerella spp. to Persist at Low Levels in the Vaginal Microbiome. Microbial Ecology 82, 1048–1060. 10.1007/s00248-020-01643-1.

84. Salliss, M.E., Maarsingh, J.D., Garza, C., Łaniewski, P., and Herbst-Kralovetz, M.M. (2021). Veillonellaceae family members uniquely alter the cervical metabolic microenvironment in a human three-dimensional epithelial model. npj Biofilms and Microbiomes 7, 57. 10.1038/s41522-021-00229-0.

85. Elwood, C., Albert, A., McClymont, E., Wagner, E., Mahal, D., Devakandan, K., Quiqley, B., Pakzad, Z., Yudin, M., Hill, J., et al. (2020). Different and diverse anaerobic microbiota were seen in women living with HIV with unsuppressed HIV viral load and in women with recurrent bacterial vaginosis: a cohort study. BJOG: An International Journal of Obstetrics & Gynaecology 127, 250–259. 10.1111/1471-0528.15930.

86. Plummer, E.L., Vodstrcil, L.A., Doyle, M., Danielewski, J.A., Murray, G.L., Fehler, G., Fairley, C.K., Bulach, D.M., Garland, S.M., Chow, E.P.F., et al. (2021). A Prospective, Open-Label Pilot Study of Concurrent Male Partner Treatment for Bacterial Vaginosis. mBio 12, 21. 10.1128/mbio.02323-21.

87. Glascock, A.L., Jimenez, N.R., Boundy, S., Koparde, V.N., Brooks, J.P., Edwards, D.J., Strauss III, J.F., Jefferson, K.K., Serrano, M.G., Buck, G.A., and Fettweis, J.M. (2021). Unique roles of vaginal Megasphaera phylotypes in reproductive health. Microbial Genomics 7. 10.1099/mgen.0.000526.

88. Rasmussen, M. (2016). Aerococcus: an increasingly acknowledged human pathogen. Clinical Microbiology and Infection 22, 22–27. 10.1016/j.cmi.2015.09.026.

89. Hočevar, K., Maver, A., Vidmar Šimic, M., Hodžić, A., Haslberger, A., Premru Seršen, T., and Peterlin, B. (2019). Vaginal Microbiome Signature Is Associated With Spontaneous Preterm Delivery. Frontiers in Medicine 6. 10.3389/fmed.2019.00201.

90. Chu, D.M., Seferovic, M., Pace, R.M., and Aagaard, K.M. (2018). The microbiome in preterm birth. Best Practice & Research Clinical Obstetrics & Gynaecology 52, 103–113. 10.1016/j.bpobgyn.2018.03.006.

91. Blostein, F., Gelaye, B., Sanchez, S.E., Williams, M.A., and Foxman, B. (2020). Vaginal microbiome diversity and preterm birth: results of a nested case–control study in Peru. Annals of Epidemiology 41, 28–34. 10.1016/j.annepidem.2019.11.004.

92. Schubert, M., Lindgreen, S., and Orlando, L. (2016). AdapterRemoval v2: rapid adapter trimming, identification, and read merging. BMC Research Notes 9, 88. 10.1186/s13104-016-1900-2.

93. MEEPTOOLS: a maximum expected error based FASTQ read filtering and trimming toolkit. (2017). International Journal of Computational Biology and Drug Design 10, 237–247. 10.1504/ijcbdd.2017.085409.

94. Schoch, C.L., Ciufo, S., Domrachev, M., Hotton, C.L., Kannan, S., Khovanskaya, R., Leipe, D., Mcveigh, R., O’Neill, K., Robbertse, B., et al. (2020). NCBI Taxonomy: a comprehensive update on curation, resources and tools. Database 2020. 10.1093/database/baaa062.

95. Xu, S., Dai, Z., Guo, P., Fu, X., Liu, S., Zhou, L., Tang, W., Feng, T., Chen, M., Zhan, L., et al. (2021). ggtreeExtra: Compact Visualization of Richly Annotated Phylogenetic Data. Molecular Biology and Evolution 38, 4039–4042. 10.1093/molbev/msab166.

96. Uritskiy, G.V., DiRuggiero, J., and Taylor, J. (2018). MetaWRAP—a flexible pipeline for genome-resolved metagenomic data analysis. Microbiome 6, 158. 10.1186/s40168-018-0541-1.

97. Olm, M.R., Brown, C.T., Brooks, B., and Banfield, J.F. (2017). dRep: a tool for fast and accurate genomic comparisons that enables improved genome recovery from metagenomes through de-replication. The ISME Journal 11, 2864–2868. 10.1038/ismej.2017.126.

98. Chaumeil, P.-A., Mussig, A.J., Hugenholtz, P., and Parks, D.H. (2022). GTDB-Tk v2: memory friendly classification with the genome taxonomy database. Bioinformatics 38, 5315–5316. 10.1093/bioinformatics/btac672.

99. Minh, B.Q., Schmidt, H.A., Chernomor, O., Schrempf, D., Woodhams, M.D., von Haeseler, A., and Lanfear, R. (2020). IQ-TREE 2: New Models and Efficient Methods for Phylogenetic Inference in the Genomic Era. Molecular Biology and Evolution 37, 1530–1534. 10.1093/molbev/msaa015.

100. Kalyaanamoorthy, S., Minh, B.Q., Wong, T.K.F., von Haeseler, A., and Jermiin, L.S. (2017). ModelFinder: fast model selection for accurate phylogenetic estimates. Nature Methods 14, 587–589. 10.1038/nmeth.4285.

101. Brooks, J.P., Buck, G.A., Chen, G., Diao, L., Edwards, D.J., Fettweis, J.M., Huzurbazar, S., Rakitin, A., Satten, G.A., Smirnova, E., et al. (2017). Changes in vaginal community state types reflect major shifts in the microbiome. Microbial Ecology in Health and Disease 28, 1303265. 10.1080/16512235.2017.1303265.

102. Rice, P., Longden, I., and Bleasby, A. (2000). EMBOSS: The European Molecular Biology Open Software Suite. Trends in Genetics 16, 276–277. 10.1016/S0168-9525(00)02024-2.

103. Hyatt, D., Chen, G.-L., LoCascio, P.F., Land, M.L., Larimer, F.W., and Hauser, L.J. (2010). Prodigal: prokaryotic gene recognition and translation initiation site identification. BMC Bioinformatics 11, 119. 10.1186/1471-2105-11-119.

104. Cantalapiedra, C.P., Hernández-Plaza, A., Letunic, I., Bork, P., and Huerta-Cepas, J. (2021). eggNOG-mapper v2: Functional Annotation, Orthology Assignments, and Domain Prediction at the Metagenomic Scale. Molecular Biology and Evolution 38, 5825–5829. 10.1093/molbev/msab293.

105. Huerta-Cepas, J., Szklarczyk, D., Heller, D., Hernández-Plaza, A., Forslund, S.K., Cook, H., Mende, D.R., Letunic, I., Rattei, T., Jensen, Lars J., et al. (2018). eggNOG 5.0: a hierarchical, functionally and phylogenetically annotated orthology resource based on 5090 organisms and 2502 viruses. Nucleic Acids Research 47, D309–D314. 10.1093/nar/gky1085.

106. Sjöqvist, C., Delgado, L.F., Alneberg, J., and Andersson, A.F. (2021). Ecologically coherent population structure of uncultivated bacterioplankton. The ISME Journal 15, 3034–3049. 10.1038/s41396-021-00985-z.

107. Mistry, J., Finn, R.D., Eddy, S.R., Bateman, A., and Punta, M. (2013). Challenges in homology search: HMMER3 and convergent evolution of coiled-coil regions. Nucleic Acids Research 41, e121–e121. 10.1093/nar/gkt263.

108. Mistry, J., Chuguransky, S., Williams, L., Qureshi, M., Salazar, Gustavo A., Sonnhammer, E.L.L., Tosatto, S.C.E., Paladin, L., Raj, S., Richardson, L.J., et al. (2020). Pfam: The protein families database in 2021. Nucleic Acids Research 49, D412–D419. 10.1093/nar/gkaa913.

109. Papadopoulos, J.S., and Agarwala, R. (2007). COBALT: constraint-based alignment tool for multiple protein sequences. Bioinformatics 23, 1073–1079. 10.1093/bioinformatics/btm076.

110. Grant, B.J., Rodrigues, A.P.C., ElSawy, K.M., McCammon, J.A., and Caves, L.S.D. (2006). Bio3d: an R package for the comparative analysis of protein structures. Bioinformatics 22, 2695–2696. 10.1093/bioinformatics/btl461.

111. Ou, J., and Zhu, L.J. (2019). trackViewer: a Bioconductor package for interactive and integrative visualization of multi-omics data. Nature Methods 16, 453–454. 10.1038/s41592-019-0430-y.

112. Gu, Z., Gu, L., Eils, R., Schlesner, M., and Brors, B. (2014). circlize implements and enhances circular visualization in R. Bioinformatics 30, 2811–2812. 10.1093/bioinformatics/btu393.

