## Supplemental Information for "Population genetic analyses of longitudinal vaginal microbiome reveal racioethnic evolutionary dynamics and prevailing positive selection of *Lactobacillus* adhesins"

### Supplementary Figures & Legends

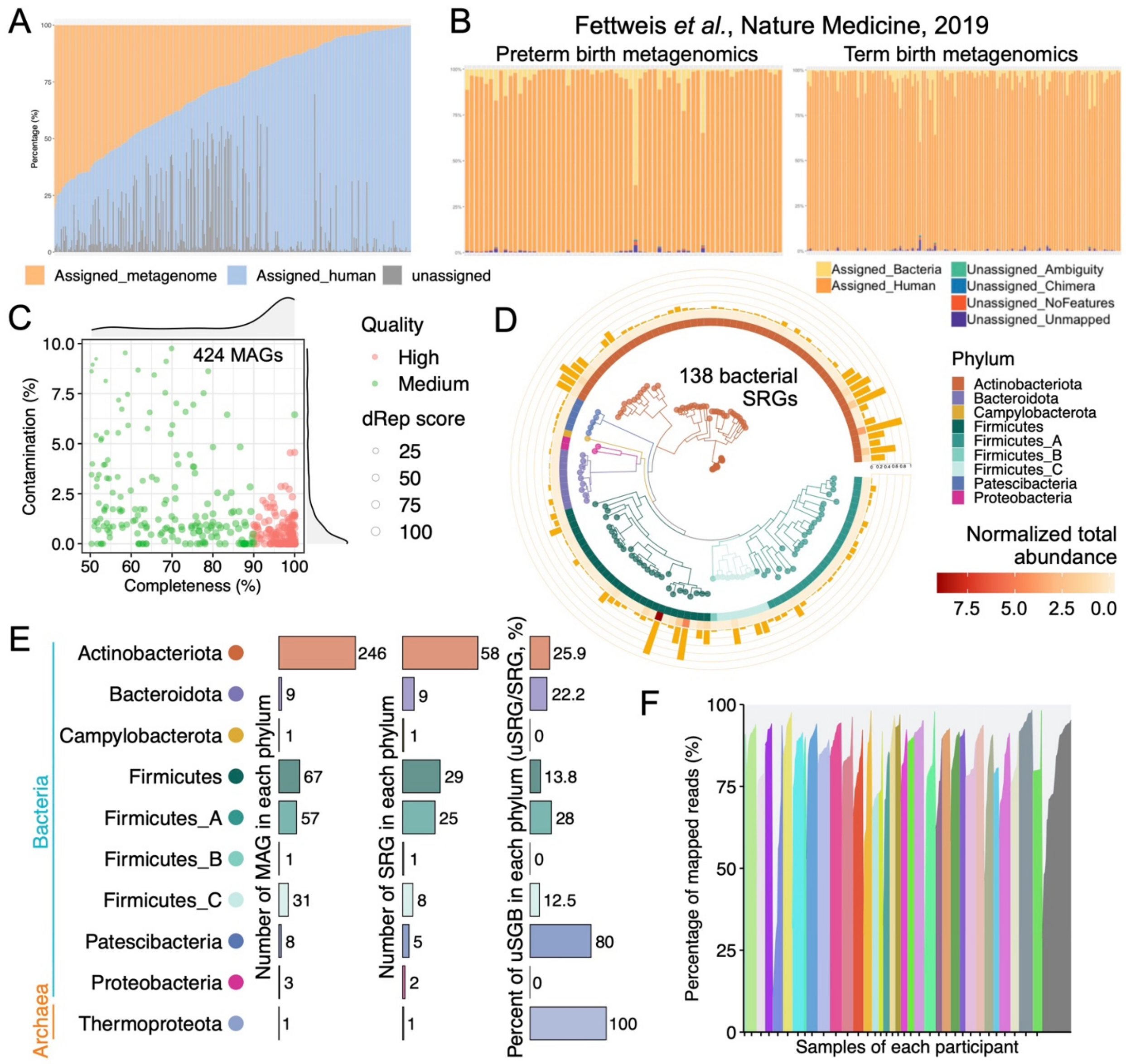


**Figure S1**

**Figure S1. Quality, taxonomy, and representatives of metagenome-assembled genomes (MAGs)**

(**A**) Assignment of metagenomics quality-filtered reads. Values are shown as the percentage abundance.

(**B**) Assignment of metagenomics quality-filtered reads in Fettweis *et al.*'s study [1]. Please refer to Extended Data Figure 4 of Fettweis *et al.*, Nature Medicine, 2019, for original figures and legends.

(**C**) The quality of 424 MAGs after removing redundancy at 98% average nucleotide identity (ANI). 240 MAGs satisfied the high-quality criteria (≥50% completeness and <10% contamination), and 184 MAGs satisfied the high-medium criteria (>90% completeness and <5% contamination [2]. The 'dRep score' was calculated by dRep software with the formula: score = (1*completeness) - (5*contamination) + (0.5*log10(ctg_N50)) + (1*log10(contig_bp)) + (2*(centrality-0.95)*100) [3]. Density plots along the x and y axes show the distribution of completeness and contamination, respectively.

(**D**) Phylogenetic tree of 138 bacterial species-level representative genomes (SRGs). The branches and inner strip were colored by phyla. The outer strip represents the normalized total abundance of each SRG in 351 samples. The orange bar shows the prevalence of each SRG.

(**E**) The number of MAGs and SGBs and the percentage of unknown SGB (uSGB) in each phylum. Only one SGB belongs to an archaeal phylum, and the others belong to bacterial phyla. The SGBs without an existing reference genome (which could not be annotated at the species level by GTDB-tk) were defined as uSGBs.

(**F**) The percentage of reads mapped to 139 SRGs. The bars of samples from the same participant have the same color.

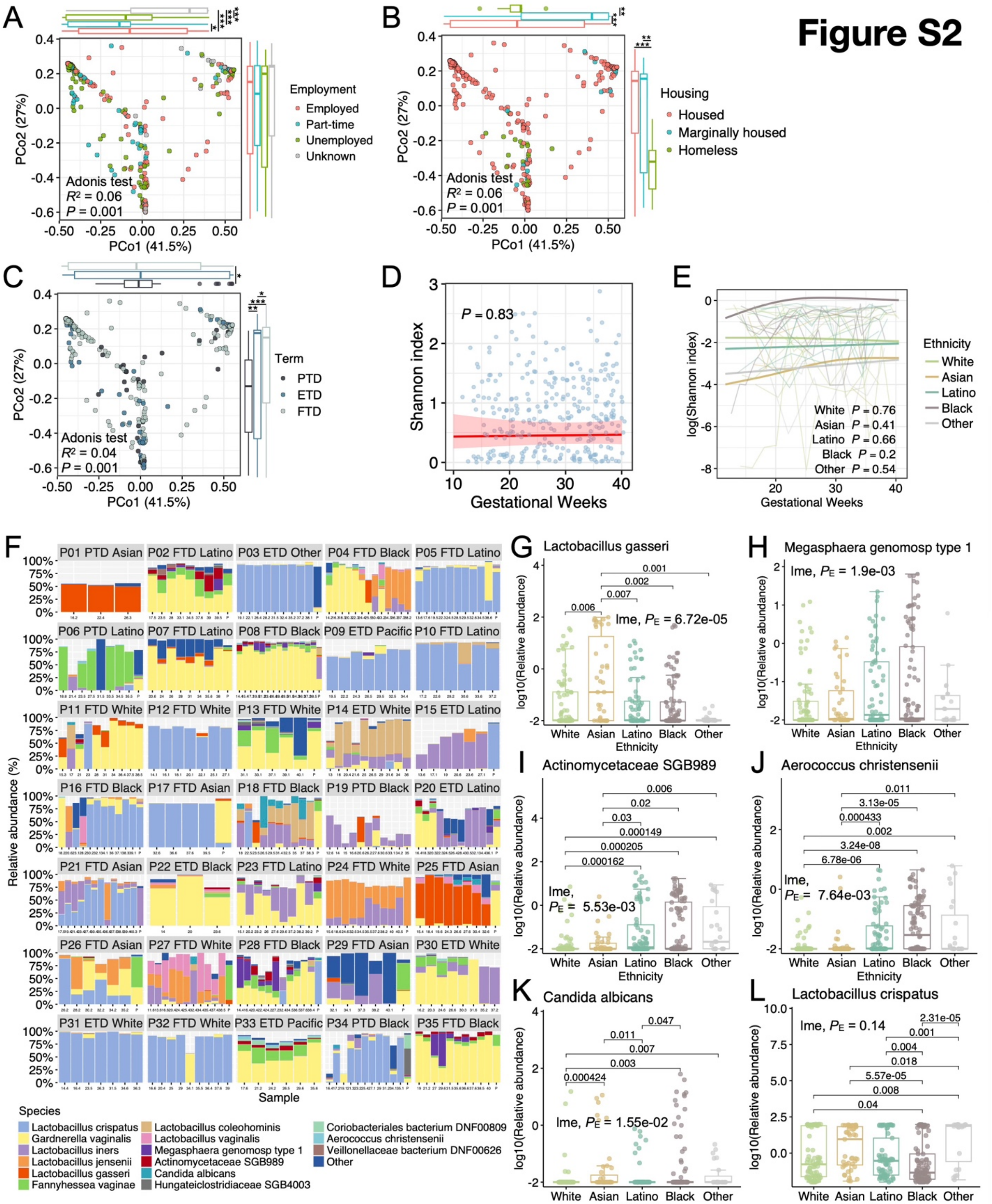
Figure S2
A
Employment
Employed
Part-time
Unemployed
Unknown
Adonis test
$R^2$ = 0.06
$P$ = 0.001
PCo1 (41.5%)
PCo2 (27%)
B
Housing
Housed
Marginally housed
Homeless
Adonis test
$R^2$ = 0.06
$P$ = 0.001
C
Term
PTD
ETD
FTD
Adonis test
$R^2$ = 0.04
$P$ = 0.001
D
$P$ = 0.83
Shannon index
Gestational Weeks
E
Ethnicity
White
Asian
Latino
Black
Other
White $P$ = 0.76
Asian $P$ = 0.41
Latino $P$ = 0.66
Black $P$ = 0.2
Other $P$ = 0.54
log(Shannon index)
Gestational Weeks
F
P01 PTD Asian
P02 FTD Latino
P03 ETD Other
P04 FTD Black
P05 FTD Latino
P06 PTD Latino
P07 FTD Latino
P08 FTD Black
P09 ETD Pacific
P10 FTD Latino
P11 FTD White
P12 FTD White
P13 FTD White
P14 ETD White
P15 ETD Latino
P16 FTD Black
P17 FTD Asian
P18 FTD Black
P19 PTD Black
P20 ETD Latino
P21 FTD Asian
P22 ETD Black
P23 FTD Latino
P24 FTD White
P25 FTD Asian
P26 FTD Asian
P27 FTD White
P28 FTD Black
P29 FTD Asian
P30 ETD White
P31 ETD White
P32 FTD White
P33 ETD Pacific
P34 PTD Black
P35 FTD Black
Relative abundance (%)
Sample
Species
Lactobacillus crispatus
Gardnerella vaginalis
Lactobacillus iners
Lactobacillus jensenii
Lactobacillus gasseri
Fannyhessea vaginae
Lactobacillus coleohominis
Lactobacillus vaginalis
Megasphaera genomosp type 1
Actinomycetaceae SGB989
Candida albicans
Hungateiclostridiaceae SGB4003
Coriobacteriales bacterium DNF00809
Aerococcus christensenii
Veillonellaceae bacterium DNF00626
Other
G
Lactobacillus gasseri
lme, $P_E$ = 6.72e-05
H
Megasphaera genomosp type 1
lme, $P_E$ = 1.9e-03
I
Actinomycetaceae SGB989
lme, $P_E$ = 5.53e-03
J
Aerococcus christensenii
lme, $P_E$ = 7.64e-03
K
Candida albicans
lme, $P_E$ = 1.55e-02
L
Lactobacillus crispatus
lme, $P_E$ = 0.14
log10(Relative abundance)
White Asian Latino Black Other
Ethnicity

27

**Figure S2. Relationships of ethnicity and other host factors to the relative abundance of dominant species in the vaginal microbiome**

(**A-C**) The principal coordinates analysis (PCoA) shows patterns of the vaginal microbiome across populations with different employment, housing statuses, and pregnancy outcomes (term). *P* indicates the *P* values of the three variables in the multivariate Adonis test. Principal components were compared using the Wilcoxon test. **P* value < 0.05; ***P* value < 0.01; ****P* value < 0.001. PTD, preterm delivery; ETD, early-term delivery; FTD, full-term delivery.

(**D**) Alpha diversity of vaginal microbiome during pregnancy. The fitted line with a 95% confidence interval represents the regression of the univariate LME model with the subject (participant) as a random effect.

(**E**) Alpha diversity of vaginal microbiome for each ethnicity during pregnancy. The fitted lines were the regression of the multivariate generalized additive mixed model (GAMM). The model incorporates ethnicity, employment, housing status, term, BMI, marriage, age, FOB, depression, a smoother gestational week, and a random subject effect to longitudinally model log-transformed alpha diversity of the vaginal microbiome.

(**F**) Vaginal microbiome profiles of all 315 samples from 35 pregnant women.

(**G-K**) Inter-ethnicity comparisons of the relative abundance of species that were significantly associated with ethnicity in Figure 2C.

(**L**) Inter-ethnicity comparisons of the relative abundance of *Lactobacillus crispatus*.

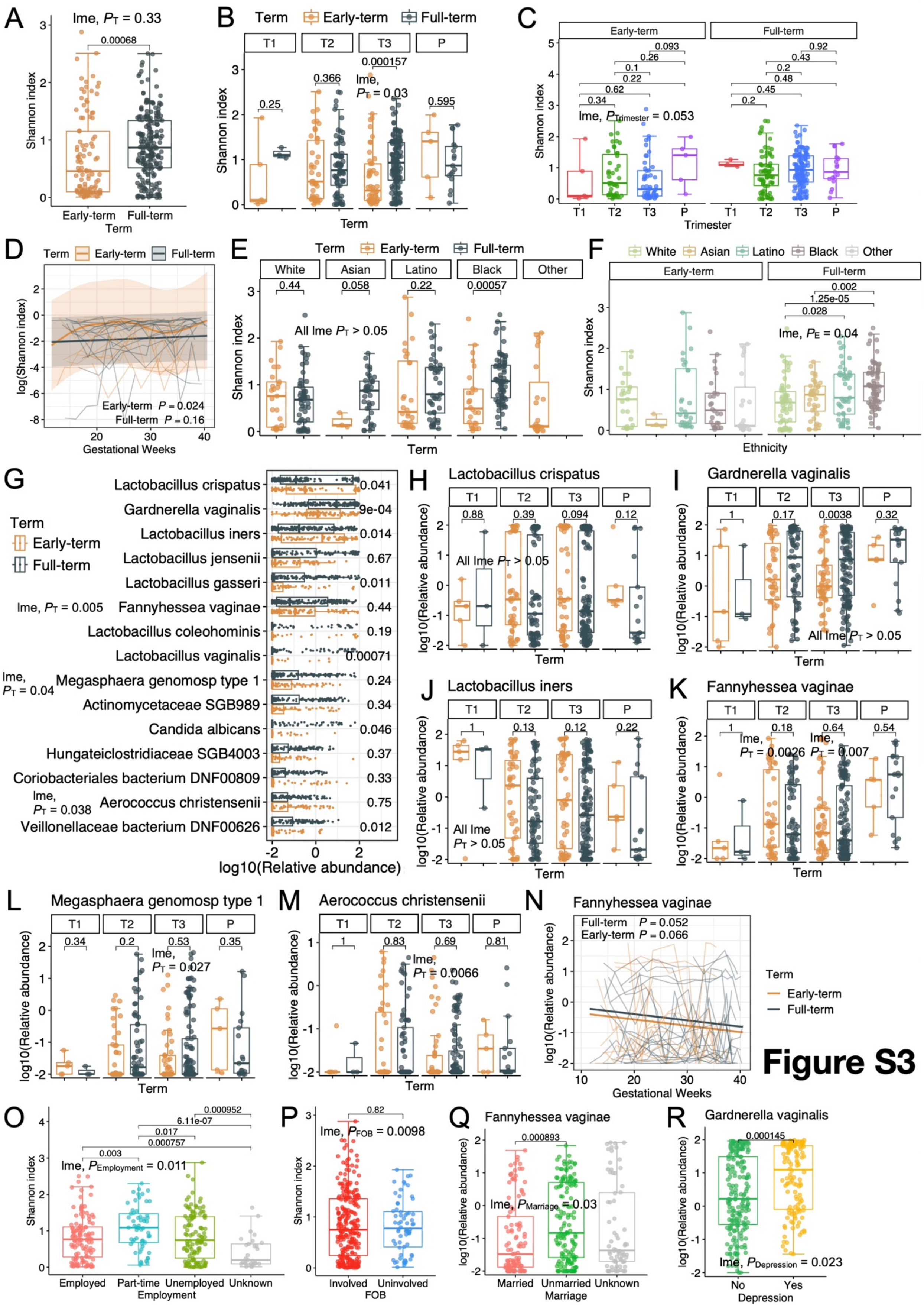
A
lme, $P_T$ = 0.33
Shannon index
Early-term
Full-term
Term
B
Term
Early-term
Full-term
T1
T2
T3
P
C
Early-term
Full-term
lme, $P_{Trimester}$ = 0.053
Trimester
D
log(Shannon index)
Early-term $P$ = 0.024
Full-term $P$ = 0.16
Gestational Weeks
E
White
Asian
Latino
Black
Other
All lme $P_T$ > 0.05
F
lme, $P_E$ = 0.04
Ethnicity
G
Lactobacillus crispatus
Gardnerella vaginalis
Lactobacillus iners
Lactobacillus jensenii
Lactobacillus gasseri
Fannyhessea vaginae
Lactobacillus coleohominis
Lactobacillus vaginalis
Megasphaera genomosp type 1
Actinomycetaceae SGB989
Candida albicans
Hungateiclostridiaceae SGB4003
Coriobacteriales bacterium DNF00809
Aerococcus christensenii
Veillonellaceae bacterium DNF00626
lme, $P_T$ = 0.005
lme, $P_T$ = 0.04
lme, $P_T$ = 0.038
log10(Relative abundance)
H
Lactobacillus crispatus
All lme $P_T$ > 0.05
I
Gardnerella vaginalis
All lme $P_T$ > 0.05
J
Lactobacillus iners
All lme $P_T$ > 0.05
K
Fannyhessea vaginae
lme, $P_T$ = 0.0026
lme, $P_T$ = 0.007
L
Megasphaera genomosp type 1
lme, $P_T$ = 0.027
M
Aerococcus christensenii
lme, $P_T$ = 0.0066
N
Fannyhessea vaginae
Full-term $P$ = 0.052
Early-term $P$ = 0.066
O
lme, $P_{Employment}$ = 0.011
Employed
Part-time
Unemployed
Unknown
Employment
P
lme, $P_{FOB}$ = 0.0098
Involved
Uninvolved
FOB
Q
Fannyhessea vaginae
lme, $P_{Marriage}$ = 0.03
Married
Unmarried
Unknown
Marriage
R
Gardnerella vaginalis
lme, $P_{Depression}$ = 0.023
No
Yes
Depression

Figure S3

**Figure S3. Vaginal microbiome differences between early-term and full-term pregnancies**

(**A**) Comparison of alpha diversity between early-term and full-term groups. The early-term group includes PTD (N = 4) and ETD (N = 9) women. *P* values of significant pair-wise comparisons were shown above the brackets. The 'lme, $P_{\mathrm{T}}$' indicates the *P* value of term (early-term, full-term) in the multivariate linear mixed-effects (LME) model.

(**B**) Comparison of alpha diversity between early-term and full-term groups in each trimester. T1, the first trimester; T2, the second trimester; T3, the third trimester.

(**C**) Alpha diversity of vaginal microbiome for women with early-term (PTD and ETD, N = 13) and full-term delivery during pregnancy.

(**D**) Comparisons of alpha diversity among trimesters for early-term and full-term groups. The 'lme, $P_{\mathrm{Trimester}}$' indicates the *P* value of the trimester in the multivariate LME model.

(**E**) Comparison of alpha diversity between early-term and full-term groups for each ethnicity.

(**F**) Comparisons of alpha diversity among ethnicities for early-term (N = 13) and full-term (N = 22) groups. The 'lme, $P_{\mathrm{E}}$' indicates the *P* value of ethnicity in the multivariate linear mixed-effects (LME) model.

(**G**) Comparisons of the relative abundance of dominant species between the early-term (N = 13) and full-term (N = 22) groups.

(**H**-**J**) The relative abundances of the three most prevalent species were compared between the early-term and full-term groups for each trimester.

(**K**-**M**) The relative abundances of species that had significant associations (lme, $P_{\mathrm{T}} < 0.05$) with the term variable in (H) were compared between the early-term and full-term groups for each trimester.

(**N**) Longitudinal multivariate GAMM of the relative abundance of *Fannyhessea vaginae* during pregnancy.

(**O**) Comparison of alpha diversity among populations with different employment statuses.

(**P**) Comparison of alpha diversity between populations with and without the involvement of the father of the baby (FOB) during pregnancy.

(**Q**) Comparison of the relative abundance of *Fannyhessea vaginae* among populations with different marriage statuses.

(**R**) Comparisons of the relative abundance of *Gardnerella vaginalis* between populations with and without depression.

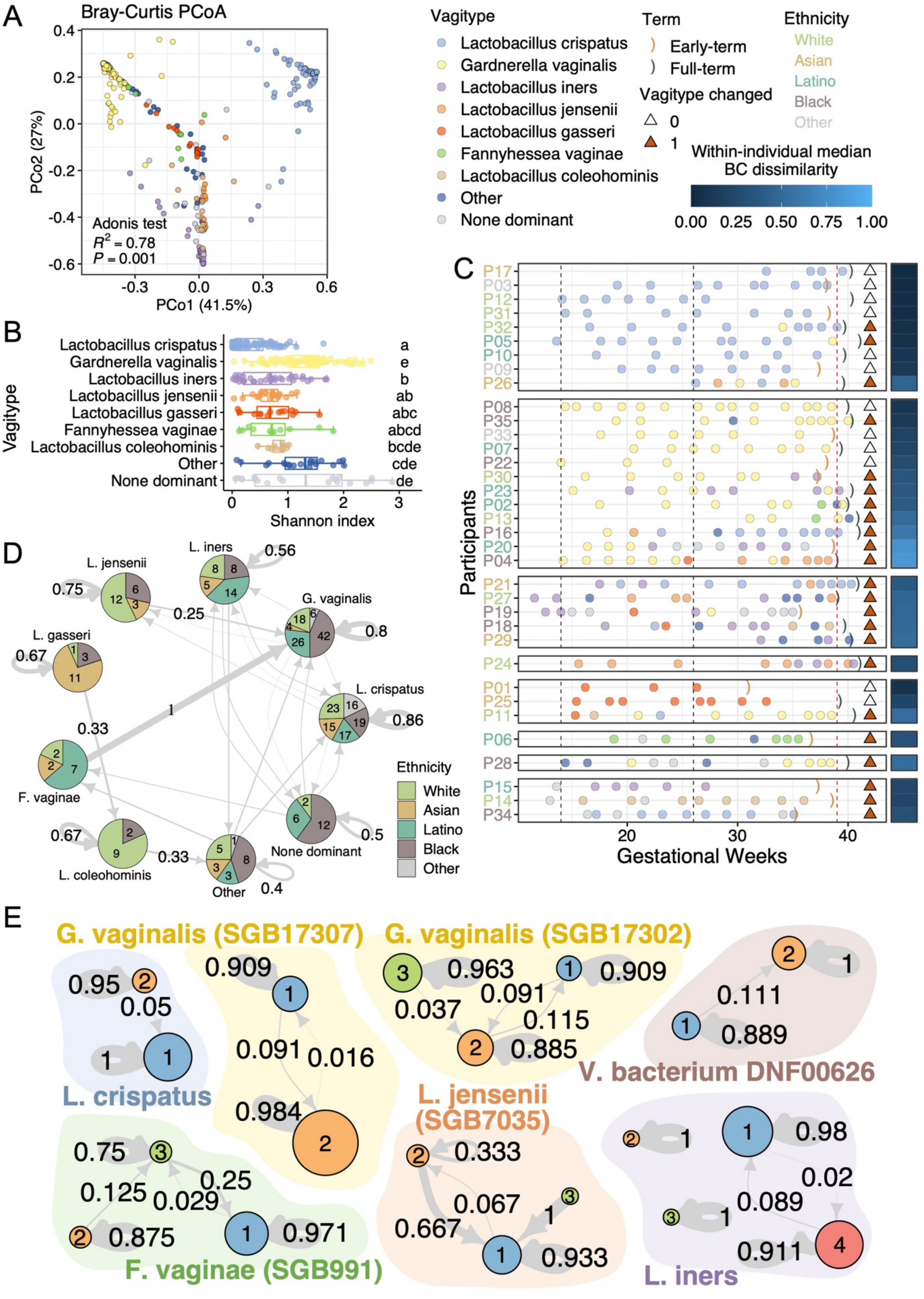
A
Bray-Curtis PCoA
PCo1 (41.5%)
PCo2 (27%)
Adonis test
$R^2$ = 0.78
$P$ = 0.001
Vagitype
Lactobacillus crispatus
Gardnerella vaginalis
Lactobacillus iners
Lactobacillus jensenii
Lactobacillus gasseri
Fannyhessea vaginae
Lactobacillus coleohominis
Other
None dominant
Term
Early-term
Full-term
Vagitype changed
Ethnicity
White
Asian
Latino
Black
Other
Within-individual median BC dissimilarity
B
Shannon index
C
Participants
Gestational Weeks
D
E
G. vaginalis (SGB17307)
G. vaginalis (SGB17302)
L. crispatus
V. bacterium DNF00626
L. jensenii (SGB7035)
F. vaginae (SGB991)
L. iners

Figure S4

**Figure S4. The transition of vagitypes and dominant strains during pregnancy**

(**A**) PCoA analysis shows distinct patterns among vagitypes. Vagitype 'none dominant' includes samples that none of the species has a relative abundance greater than 30%, possibly due to a high unclassified ratio or having multiple species with an abundance lower than 30%. The 'other' group includes vagitypes defined in no more than five samples.

(**B**) Comparison of alpha diversity among vagitypes. The letters on the right refer to the results of Tukey's test of all pairwise comparisons. Different letters indicate statistical significance.

(**C**) Longitudinal transitions of vagitypes in this study. The triangles on the right indicate whether vagitypes changed during pregnancy (filled brown). The vertical dashed lines and brackets are described in Figure 1A. Participants were divided into several panels according to the vagitypes of their first samples and were ordered by the median of Bray-Crutis (BC) dissimilarity within individuals. The colors of PIDs indicate ethnicity.

(**D**) Transition network of the vagitypes during pregnancy depicted as a Markov chain. The transition probabilities greater than 0.25 are shown on the edges (see all probabilities in Table S10). The ethnic composition of each vagitype was represented as a pie. The numbers on the pie plots indicate the number of samples.

(**E**) Strain transitions of several reference species or subspecies in the vaginal microbiome during pregnancy. Reference subspecies with the same species name are tagged in brackets with the genome ID (SGB+numbers). The nodes represent strain clusters (1, blue; 2, orange; 3, green; 4, red) that were defined by hierarchical clustering of SNP haplotype dissimilarity (see Methods).

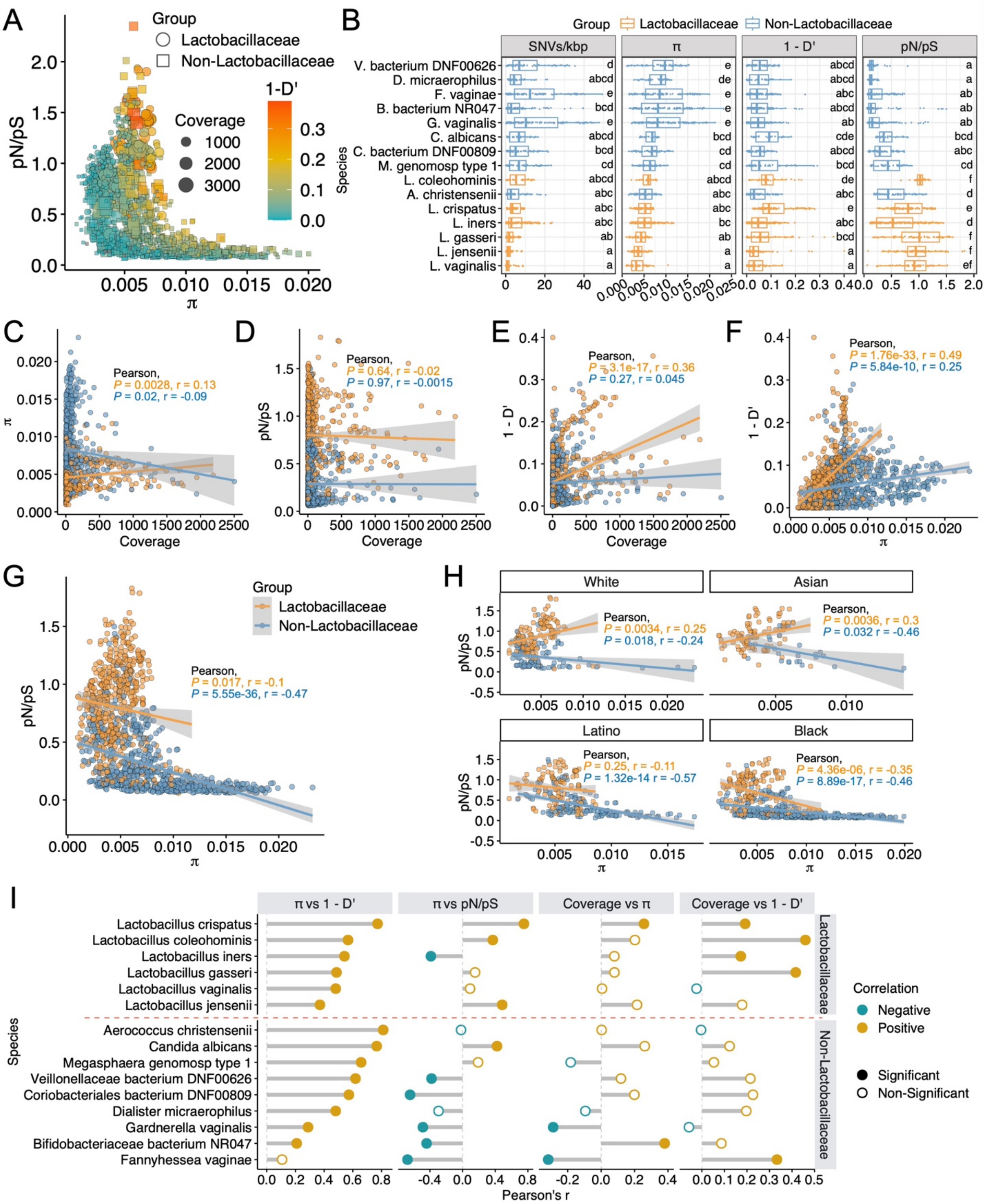


103

**Figure S5**

**Figure S5. The distributions and relationships of evolutionary metrics for vaginal microbiome**

(**A**) Overview of the evolutionary metrics of 15 species (used MAGs as references instead of the NCBI reference genomes) across all samples. Similar to Figure 4A. Each point or square represents a MAG of certain species, totaling 1613 genomes in all samples (Table S12).

(**B**) The distributions of evolutionary metrics for each species. The letters on the right refer to the results of Tukey's test of all pairwise comparisons. Different letters indicate statistical significance.

(**C**-**E**) The relationships between coverage and π, pN/pS, and 1-D' of *Lactobacillaceae* and non-*Lactobacillaceae* groups. The fitted lines with 95% confidence intervals were the regressions of linear models.

(**F**-**G**) The relationships between π and 1-D' and pN/pS.

(**H**) The relationships between π and pN/pS for each ethnicity.

(**I**) The relationships among evolutionary metrics and between coverage and π and 1-D' for each species.

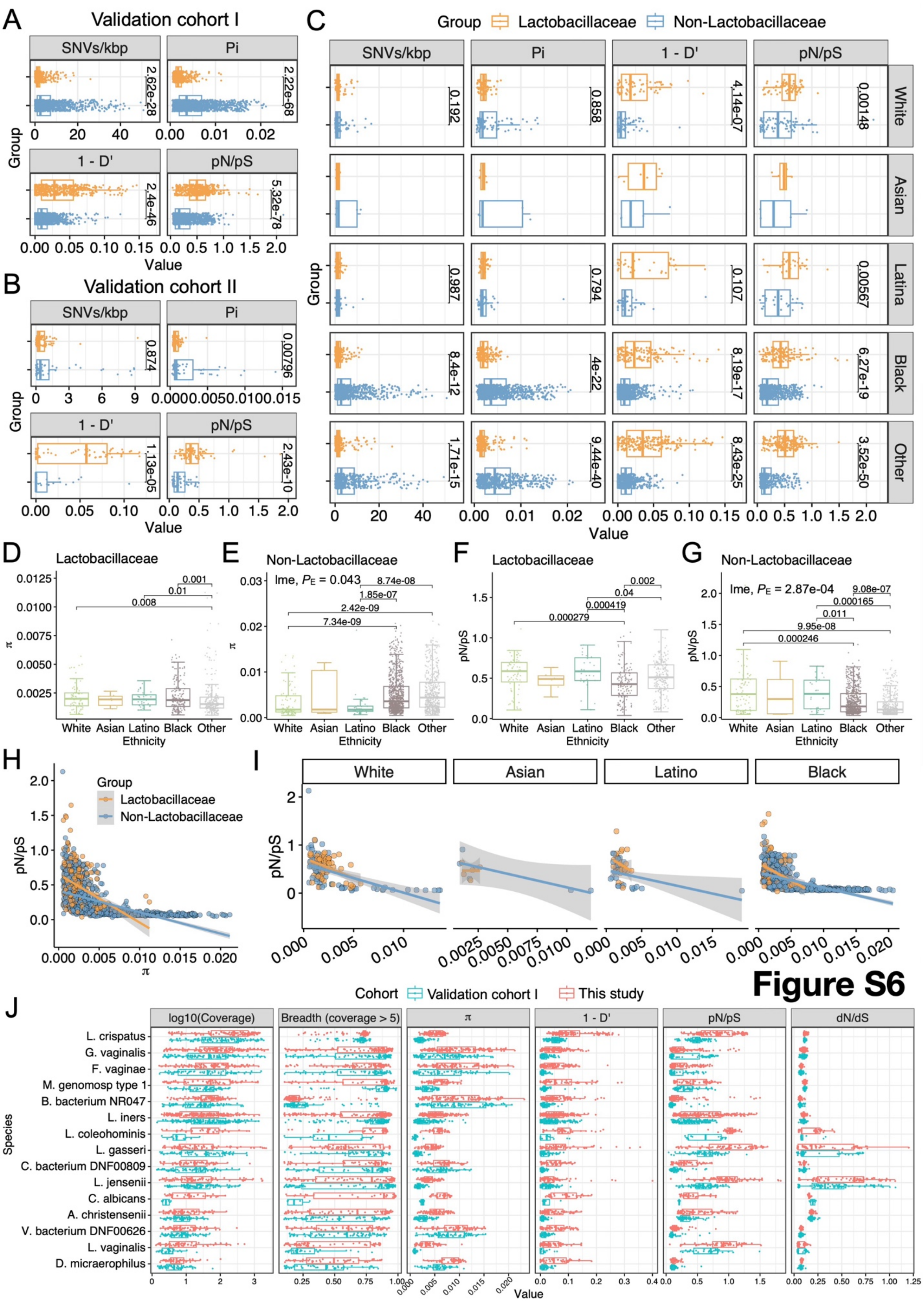


Figure S6

119

**Figure S6. Validation of the distinct evolutionary dynamics between *Lactobacillaceae* and non-*Lactobacillaceae* species and among ethnicities in independent cohorts**

(**A**) Comparisons of evolutionary metrics between *Lactobacillaceae* (orange) and non-*Lactobacillaceae* (blue) groups across all samples of the validation cohort I. The 36 species with sufficient coverage (≥ 5x across ≥ 10% of genome length) in more than 20 samples were included. Each point or square represents a genome of certain species, totaling 1645 genomes in all samples (Table S14). D’ is a measurement of linkage equilibrium ranging from 0 to 1. A higher 1-D' indicates a higher recombination rate. π, nucleotide diversity; pN/pS, the ratio of non-synonymous rate divided by synonymous rate.

(**B**) Similar to (A), but for validation cohort II. The 26 species with sufficient coverage (≥ 5x across ≥ 10% of genome length) in more than one sample were included.

(**C**) Comparisons of evolutionary metrics between *Lactobacillaceae* and non-*Lactobacillaceae* groups for each ethnicity in validation cohort I.

(**D**-**G**) Comparisons of π and pN/pS among ethnicities for *Lactobacillaceae* and non-*Lactobacillaceae* groups in validation cohort I.

(**H**) The relationships between π and pN/pS for *Lactobacillaceae* and non-*Lactobacillaceae* groups in validation cohort I.

(**I**) The relationships between π and pN/pS for *Lactobacillaceae* and non-*Lactobacillaceae* groups of each ethnicity in validation cohort I.

(**J**) The distributions of evolutionary dynamics, coverage, and breadth (percent of genome length covered by at least 5 reads) for each species of the samples from this study and the validation cohort I.

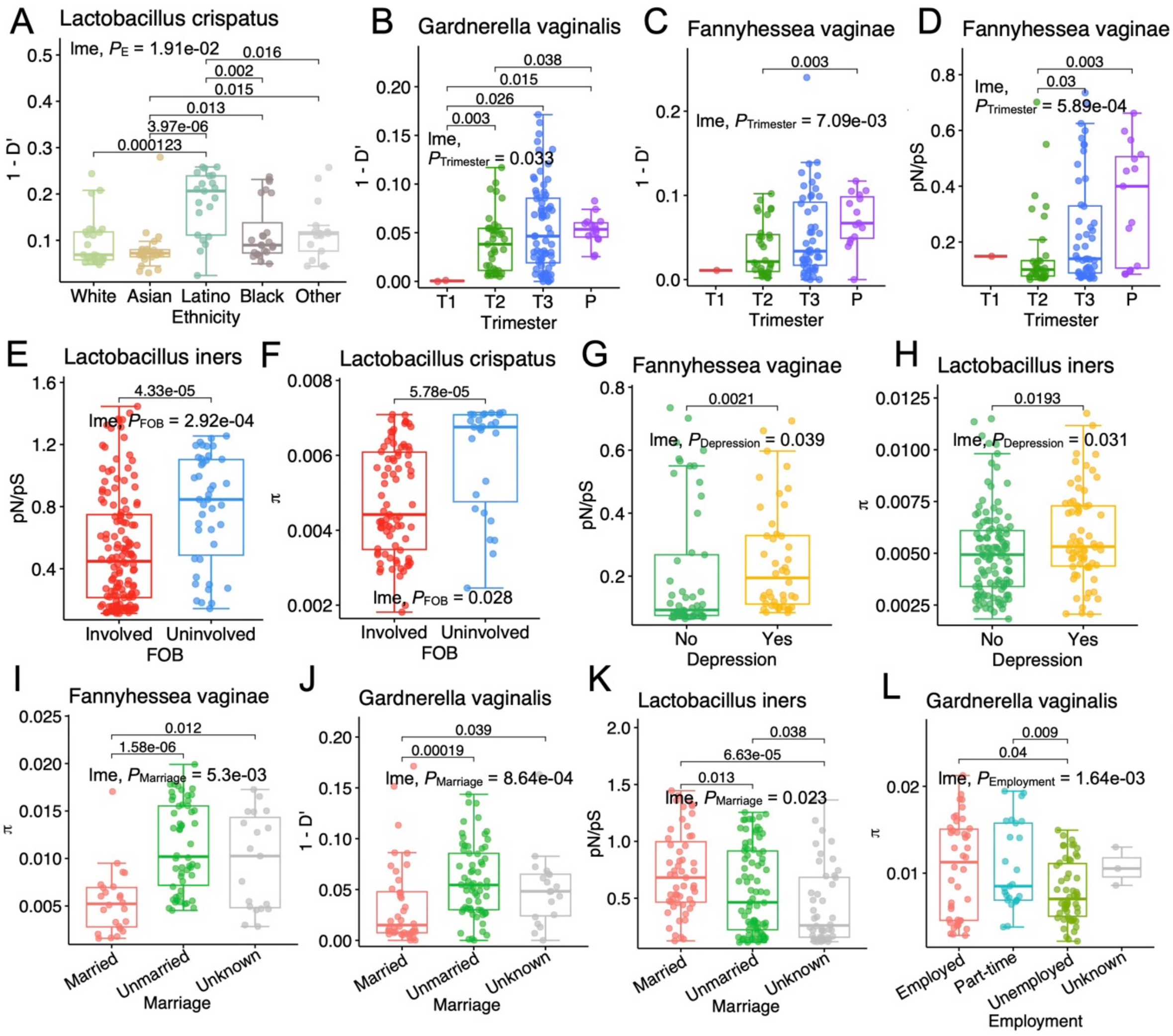


**Figure S7**

**Figure S7. The associations between evolutionary dynamics of the vaginal microbiome and FOB, depression, marriage, and employment**

(**A**) Comparisons of *Lactobacillus crispatus* recombination rate (1-D') among ethnicities.

(**B**-**D**) Changes in 1-D' and pN/pS in specific species among trimesters and postpartum.

(**E**-**F**) Comparisons of pN/pS and π of specific species between populations with and without the involvement of the father of the baby (FOB) during pregnancy.

(**G**-**H**) Comparisons of pN/pS and π of specific species between populations with and without depression.

(**I**-**K**) Comparisons of π, 1-D', and pN/pS of specific species among populations with different marriage statuses.

(**L**) Comparisons of π of *Gardnerella vaginalis* among populations with different employment statuses.

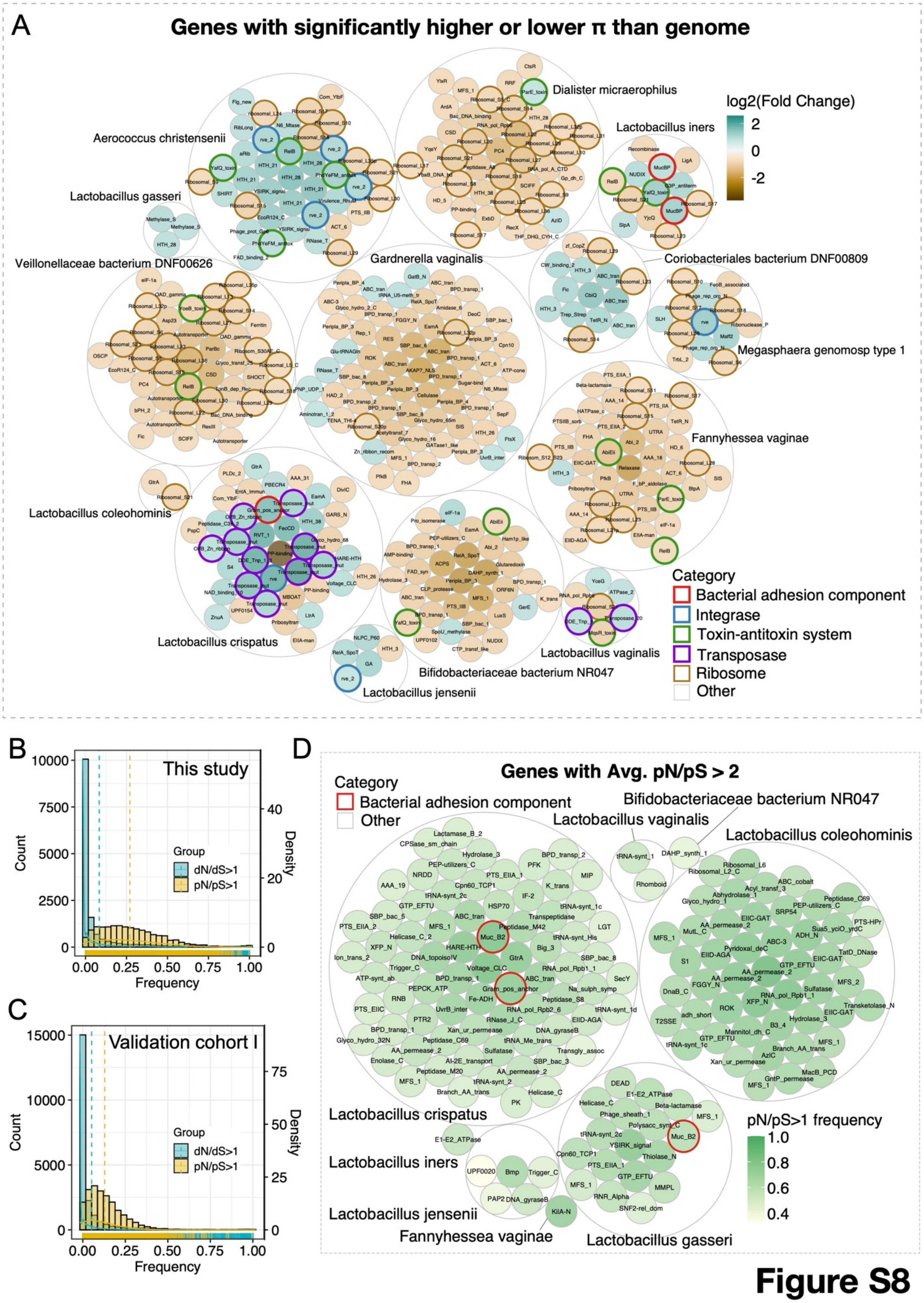


Figure S8

157

**Figure S8. The distributions of π and pN/pS of the microbial genes in the vaginal microbiome**

(**A**) The genes with significantly higher or lower π than the genomic average. Each small circle represents a gene filled based on the fold change (mean gene π / genomic average π). 353 genes of 14 species are shown. A larger gray circle packed genes of the same species. Genes of interest were marked by bold circles colored with the assigned categories.

(**B**-**C**) Distributions of the frequency of dN/dS > 1 (blue) and pN/pS > 1 (yellow) of genes with sufficient coverage (≥ 5x across ≥ 50% of gene length) in more than ten samples in this study and the validation cohort I. The vertical dashed line represents the mean frequency of each group.

(**D**) The genes with average pN/pS greater than 1. Each small circle represents a gene filled based on pN/pS > 1 frequency across samples. 149 genes of 8 species are shown. A larger gray circle packed genes of the same species. Genes of interest were marked by bold circles colored with the assigned categories.

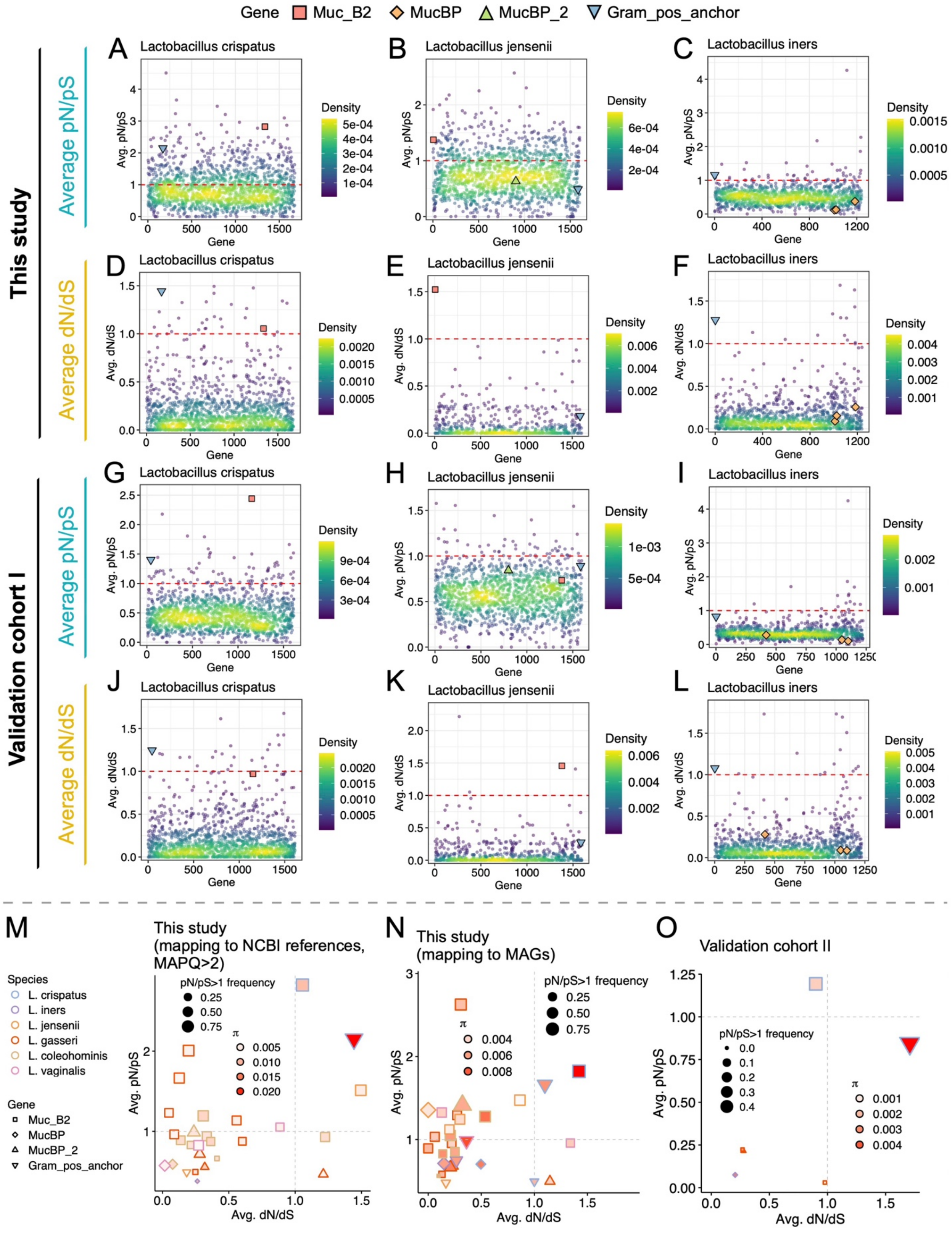
Gene
Muc_B2
MucBP
MucBP_2
Gram_pos_anchor
This study
Validation cohort I
Average pN/pS
Average dN/dS
A
Lactobacillus crispatus
B
Lactobacillus jensenii
C
Lactobacillus iners
D
Lactobacillus crispatus
E
Lactobacillus jensenii
F
Lactobacillus iners
G
Lactobacillus crispatus
H
Lactobacillus jensenii
I
Lactobacillus iners
J
Lactobacillus crispatus
K
Lactobacillus jensenii
L
Lactobacillus iners
Avg. pN/pS
Avg. dN/dS
Gene
Density
M
This study
(mapping to NCBI references, MAPQ>2)
Species
L. crispatus
L. iners
L. jensenii
L. gasseri
L. coleohominis
L. vaginalis
Gene
Muc_B2
MucBP
MucBP_2
Gram_pos_anchor
pN/pS>1 frequency
π
N
This study
(mapping to MAGs)
O
Validation cohort II
Avg. pN/pS
Avg. dN/dS

**Figure S9**

**Figure S9. The average dN/dS and pN/pS of adhesion genes**

(**A-L**) The average dN/dS and pN/pS of genes across genomes of *Lactobacillus crispatus*, *Lactobacillus jensenii*, and *Lactobacillus iners* in samples of this study (A-F) and the validation cohort I (G-L). Genes were colored by the 2D density, while *Lactobacillus* adhesins were indicated by points with different shapes and colors. The red dashed lines were drawn at the average dN/dS or pN/pS equal to 1.

(**M**) The average dN/dS and pN/pS of adhesion genes in samples of this study, calculated based on mapping to NCBI reference genomes and reads with MAPQ > 2.

(**N**) The average dN/dS and pN/pS of adhesion genes in samples of this study, calculated based on mapping to SRGs (*de novo* references generated in this study).

(**O**) The average dN/dS and pN/pS of adhesion genes in samples of the validation cohort II.

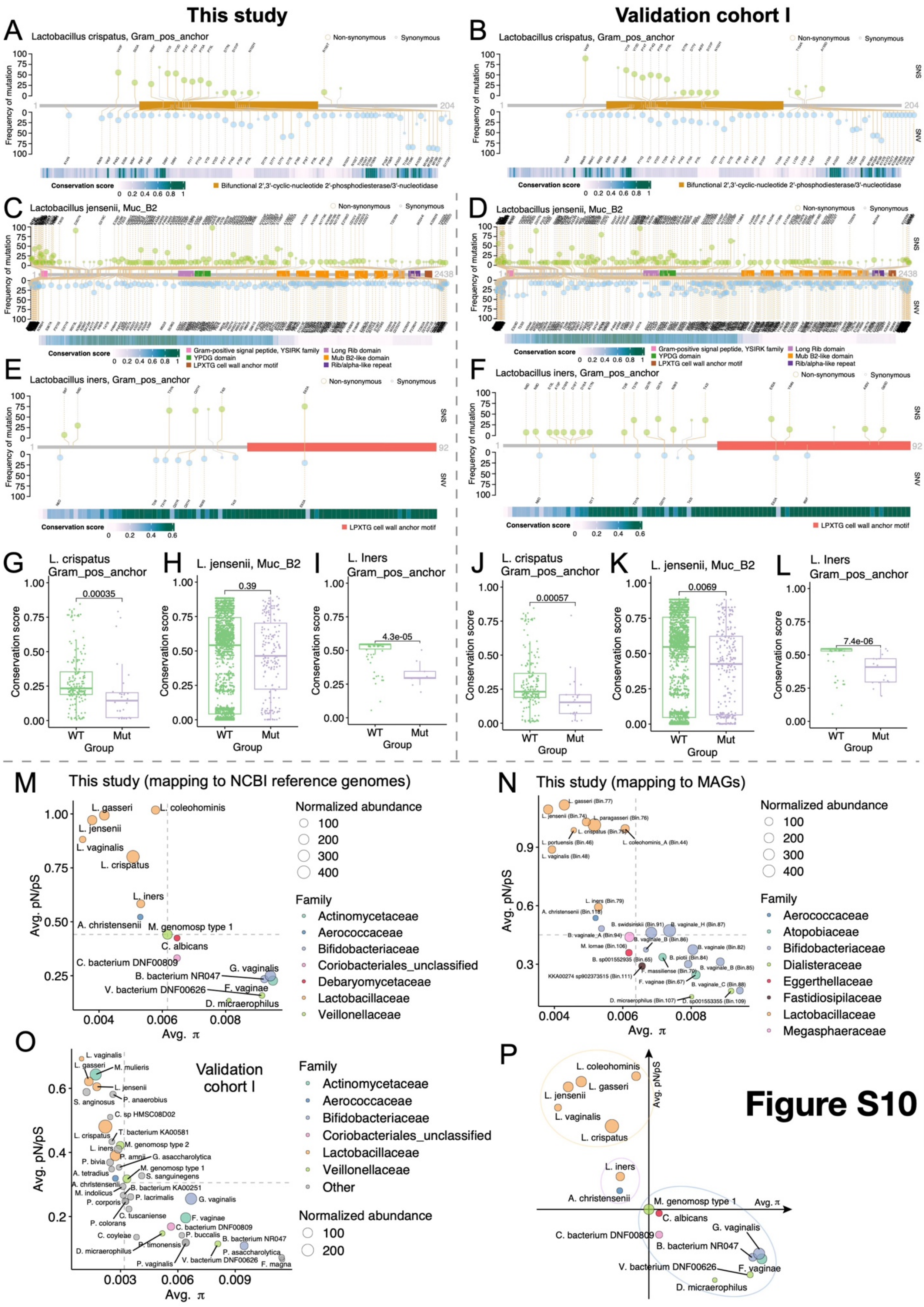
This study
Validation cohort I
A
Lactobacillus crispatus, Gram_pos_anchor
B
Lactobacillus crispatus, Gram_pos_anchor
Non-synonymous
Synonymous
Frequency of mutation
SNS
SNV
Conservation score
Bifunctional 2',3'-cyclic-nucleotide 2'-phosphodiesterase/3'-nucleotidase
C
Lactobacillus jensenii, Muc_B2
D
Lactobacillus jensenii, Muc_B2
Gram-positive signal peptide, YSIRK family
YPDG domain
LPXTG cell wall anchor motif
Long Rib domain
Mub B2-like domain
Rib/alpha-like repeat
E
Lactobacillus iners, Gram_pos_anchor
F
Lactobacillus iners, Gram_pos_anchor
LPXTG cell wall anchor motif
G
L. crispatus Gram_pos_anchor
H
L. jensenii, Muc_B2
I
L. Iners Gram_pos_anchor
J
L. crispatus Gram_pos_anchor
K
L. jensenii, Muc_B2
L
L. Iners Gram_pos_anchor
Conservation score
0.00035
0.39
4.3e-05
0.00057
0.0069
7.4e-06
WT
Mut
Group
M
This study (mapping to NCBI reference genomes)
N
This study (mapping to MAGs)
Avg. pN/pS
Avg. π
Normalized abundance
Family
Actinomycetaceae
Aerococcaceae
Bifidobacteriaceae
Coriobacteriales_unclassified
Debaryomycetaceae
Lactobacillaceae
Veillonellaceae
Atopobiaceae
Dialisteraceae
Eggerthellaceae
Fastidiosipilaceae
Megasphaeraceae
O
Validation cohort I
Other
P
Figure S10

184

**Figure S10. The mutation hotspots and conservation score of *Lactobacillus* adhesins and the proposed model of microbial evolving space defined by microbial evolutionary metrics in vaginal environments**

(**A-F**) The mutation hotspots on the amino acid sequences of *Lactobacillus* adhesins in this study (left; A, C, and E) and the validation cohort I (right; B, D, and F). Two types of mutations, single nucleotide substitution (SNS, green) and single nucleotide variant (SNV, blue), were illustrated on the two sides of the x-axis. Non-synonymous mutations were indicated by a bigger dot than synonymous mutations. The sequences and more information about *gram_pos_anchor* (Accession ID: WP_158181626.1) in *Lactobacillus crispatus*, *muc_B2* in *Lactobacillus jensenii* (Accession ID: WP_075362409.1), and *gram_pos_anchor* in (Accession ID: QFZ99646.1) in *Lactobacillus iners* are recorded in the NCBI Protein database (https://www.ncbi.nlm.nih.gov/protein). The conservation score of each amino acid was displayed by the horizontal bar below.

(**G-L**) The comparisons of conservation scores between wild-type loci (WT) and mutated loci (Mut) in *Lactobacillus* adhesins in this study (left; G-I) and the validation cohort I (right; J-L).

(**M**) The average pN/pS and $\pi$ of species across samples of this study, calculating based on NCBI reference genomes. The dashed gray lines indicate the median of average pN/pS and $\pi$.

(**N**) Similar to (M), but calculated based on MAGs.

(**O**) Similar to (M), but for validation cohort I.

(**P**) The proposed model for evolving space for bacterial species, modified from (M).

**Supplementary Table 1**

Metadata of samples in this study.

**Supplementary Table 2**

The abundance profiles given by MetaPhlAn4.

**Supplementary Table 3**

CSTs and vagitypes assignment.

**Supplementary Table 4**

The proportion of overlapped samples between CSTs and Vagitypes.

**Supplementary Table 5**

CST distribution of the first sample of each participant among ethnic groups.

**Supplementary Table 6**

Vagitype distribution of the first sample of each participant among ethnic groups.

**Supplementary Table 7**
CST distribution of the first sample of each participant of PTD, ETD, and FTD groups.

**Supplementary Table 8**
Vagitype distribution of the first sample of each participant of PTD, ETD, and FTD groups.

**Supplementary Table 9**
CST biweekly transition probabilities.

**Supplementary Table 10**
Vagitype biweekly transition probabilities.

**Supplementary Table 11**
Evolutionary metrics of 15 species across all samples in this study.

**Supplementary Table 12**
Evolutionary metrics of 15 species across all samples in this study (used MAGs as references instead of the NCBI reference genomes).

**Supplementary Table 13**
Metadata for the validation cohort I and II.

**Supplementary Table 14**
Evolutionary metrics of 36 species across all samples of the validation cohort I.

**Supplementary Table 15**
Evolutionary metrics of genes across all samples in this study.

**Supplementary Table 16**
Evolutionary metrics of genes across all samples of the validation cohort I.

### References

1. Fettweis, J.M., Serrano, M.G., Brooks, J.P., Edwards, D.J., Girerd, P.H., Parikh, H.I., Huang, B., Arodz, T.J., Edupuganti, L., Glascock, A.L., et al. (2019). The vaginal microbiome and preterm birth. Nature Medicine *25*, 1012-1021. 10.1038/s41591-019-0450-2.
2. Bowers, R.M., Kyrpides, N.C., Stepanauskas, R., Harmon-Smith, M., Doud, D., Reddy, T.B.K., Schulz, F., Jarett, J., Rivers, A.R., Eloe-Fadrosh, E.A., et al. (2017). Minimum information about a single amplified genome (MISAG) and a metagenome-assembled genome (MIMAG) of bacteria and archaea. Nature Biotechnology *35*, 725-731. 10.1038/nbt.3893.
3. Olm, M.R., Brown, C.T., Brooks, B., and Banfield, J.F. (2017). dRep: a tool for fast and accurate genomic comparisons that enables improved genome recovery from metagenomes through de-replication. The ISME Journal *11*, 2864-2868. 10.1038/ismej.2017.126.
